# Evaluation of the appropriateness of pesticide administration-concentrations to honey bee *Apis mellifera* colony in long-term field experiments and modest proposal

**DOI:** 10.1101/2023.08.25.554728

**Authors:** Toshiro Yamada

## Abstract

For investigating the effects of pesticides on bee colonies in actual apiaries, field experiments in an open environment are effective. It is natural to assume that the total food or pesticide intake taken by a honey bee colony includes their intakes from nature in addition to each administered in the field experiment. Therefore, in field experiments, setting the administration-concentration of pesticides for honey bee colonies is one of the most difficult tasks.

The validity of pesticide concentrations administered to bee colonies was evaluated by various intakes of pesticides and foods obtained from five long-term field experiments conducted in Midwest Japan with distinct changes of four seasons and Maui, U.S.A. without cold winter.

The author found a relationship, log***P_BD_***=log***C_P_***–2.4, between pesticide concentration (***C_P_***) and the average daily intake of pesticide ingested by one bee (***P_BD_***), and found that their ratio ***P_BD_***/***C_P_*** was about 1/250, which was significantly lower than the ***LD_50_*** value.

This fact supports that the concentration of pesticide administration in long-term field experiments conducted so far was not too high, but rather reasonable.

In addition, the average daily intakes of sugar syrup and pollen paste ingested by one bee was found to be 3.1 mg/bee/day and 0.8 mg/bee/day, respectively, and notwithstanding their wide variation, be roughly constant regardless of the concentration of pesticides.

I believe that this paper will help provide new guidelines for setting pesticide dosing concentrations, which have not been clear so far, in field experiments on the administration of pesticides to bee colonies.

## Introduction

In recent years, the risk to the ecosystem of neonicotinoid pesticides (hereinafter referred to as neonics), which exhibit long-term residual effects, toxic and permeable neurotoxicity, has been pointed out (Tirado et al., 2013; Goulson, 2013; Wood & Goulson, 2017). In particular, there have been concerns about the negative impact of neonics on bees. In long-term field administration experiments of neonics to bees that have been conducted to investigate the effects of neonics on bees, it has often been debated whether the concentration of pesticides administered to a honey bee colony is actually appropriate. A median lethal dose (***LD_50_***) for bees has often been used as an indicator to assess whether the pesticide administration-concentration is appropriate.

It is well known that honeybees, which are eusocial insects, have their roles determined by caste and act on a colony basis. Castes in a colony are divided into three types: one queen bee, a small number of male bees (drones) and many female bees (worker bees).

In honeybee society, the three roles of egg-laying, reproduction and rearing are shared by the queen bee, drone and worker bee, respectively. While being cared for by worker bees, the queen bee continues to lay eggs for the rest of its life after mating, and the drone spends its entire life with mating as its only job. Worker bees are responsible for almost all other tasks such as foraging and larval rearing.

In worker bees, the division of labor is further sub-divided, and it is roughly divided into in-house bees that are responsible for cleaning and rearing in the hive box and foraging bees that collect pollen and nectar outside the hive box. After emerging, worker bees work as in-house bees in the hive box, and then, their age in day, forage outside the hive box as foraging bees after they become old.

The order of this division of labor changes depending on the state of the colony, the surrounding environment in which the colony is placed, the season, etc. However, in the case that the momentum of the colony is strong, the surrounding pollen and nectar sources are abundant, and active foraging activities are carried out, the change in division of labor corresponds to the progression of day-age (Watanabe, H. & Watanabe, T., 1974).

In addition to the peculiarities of honey bee colonies, which consist of such a complex social structure consisting of the caste system and division of labor, long-term field experiments in an environment close to real apiaries and measurement of ***LD_50_*** in the laboratory (laboratory experiments) are very different, as shown in Supplementary Table S1, which summarizes the differences between the characteristics of laboratory experiments when measuring the median lethal dose (***LD_50_***) of bees and those of long-term field experiments of honey bee colonies. The main differences between two are as follows: In long-term field experiments, it is easy to reproduce an experimental environment close to that of an actual apiary, but in laboratory experiments used for ***LD_50_*** measurement, it is extremely difficult to reproduce the experimental environment of an actual apiary. On the other hand, it is easy to control the experimental environment in laboratory experiments conducted in a closed system, but it is extremely difficult to control the experimental environment in long-term field experiments conducted in an open system.

For example, long-term field experiments are conducted in an open natural environment with a wide degree of freedom, while laboratory experiments such as the measurement of ***LD_50_*** are conducted in a closed environment with a narrow degree of freedom. In other words, at the time of ***LD_50_*** measurement, the ecological environment of honeybees is controlled, the intake and timing of food containing pesticides by bees are accurately controlled and measured, and food other than the pesticide-containing food administered in the experiment is not ingested.

On the other hand, in long-term field experiments, honey bee colonies are composed of a social structure that maintains a caste system and division of labor in almost the same way as actual apiaries, and the experiments are conducted in the same natural environment as the apiary, and the intake and timing of the administered pesticide-containing food are not controlled and are free. In addition, the consumed pesticide-containing food is often mixed with food collected from nature and stored in the hive box, and the timing and intake of such stored food are freely carried out according to the will of the honeybees and are not managed.

In conducting field experiments in an open system where it is difficult to control the experimental environment, we made the following efforts to reduce the influence of the experimental environment as much as possible (for details, see T. Yamada et al., 2012, 2018_a_, 2018_b_, 2018_c_, 2018_d_; T. Yamada & K. Yamada, 2020; Yamada, 2020): (1) A pesticide-free drinking fountain was set up in the experimental site, and pesticide-free plants and flowers and trees were planted to prevent bees from ingesting pesticides from sources other than the experiment; (2) The control colony and the experimental colony were arranged alternately to reduce the influence of the location of the hive box on the experimental results; (3) In order to reduce the errors in the intakes of sugar syrup and pollen paste, and to reduce the measurement error of the numbers of adult bees and capped brood in the hive box and the dead bees in and around the hive box, immediately after dawn before the bees start foraging activities, measure the remaining amount of sugar syrup or pollen paste, and take photos of each side of all comb frames and the five walls in the hive box (can be checked repeatedly), using newly developed counting software as an adjunct to these photographs, the number of adult bees and capped brood were calculated from these photographs. And dead bees in and around the hive box were counted one by one and their number was recorded. Bees affected by mites, while enlarging photos of comb frames and the four walls and bottom of the hive box, were counted.

By the way, if the difference in preference of honeybees is between the pesticide-containing food administered (sugar syrup, pollen paste) and the food collected from nature (flower nectar, bee bread), there is a difference between the apparent intake (including stored food) and the true intake of pesticide-containing food by honeybees, and the true pesticide intake may be inaccurate. In this way, honey bee colonies in long-term field experiments constitute a complex social structure composed of caste system and division of labor, and operate freely in an open natural environment that is not controlled. On the other hand, in the measurement environment of ***LD_50_*** in laboratory experiments, the honey bee colony consists of only a small number of worker bees without caste or division of labor, and is engaged in constrained activities in a highly controlled environment.

Thus, the appropriateness of the administration-concentration of pesticides in long-term field experiments in an open ecological environment in which honey bee colonies maintaining the caste system and division of labor as almost complete eusocial insects can be freely ingested from other than pesticide-containing food to which they are administered, from ***LD_50_*** measured in a very incomplete social structure consisting of a small number of bees and a controlled closed ecological environment, as in Nakamura et al. (2014), it seems reckless to evaluate.

For example, judging from the ***LD_50_*** value, even if food (sugar solution, pollen paste) with a pesticide concentration that seems to be too high is administered to bees in a long-term field experiment, the behavior of bees is free in the field experiment, so if the bees hardly ingest the administered pesticide-containing food and preferentially ingest food procured from the field. Alternatively, it may also happen that the actual pesticide intake of bees is lower than the ***LD_50_*** value. That is, from the ***LD_50_*** value in a closed measurement environment in which all ingested food contains pesticides, it is expected that the pesticide administration-concentration in the field experiment where bees are allowed to move freely is expected to be significantly different from the concentration of pesticide administration in a field experiment where bees are allowed to move. Similarly, in field experiments, food containing pesticides that appears to have been ingested is often stored in the cell. Since it is up to the bees to decide when and how much of the stored pesticide they actually ingest, the actual intake is even more uncertain.

Therefore, in the long-term field experiments that we have conducted so far, we estimated the pesticide intake per bee during the pesticide administration period from the total amount (intake) of food containing pesticides in each colony and the total number of adult bees (initial number + number of adult bees that have newly eclosed calculated from the number of capped brood) (T. Yamada et al., 2018_a_, 2018_b_).

In this paper, we estimate the average amount of pesticide ingested by one bee per day during the pesticide administration period in a long-term field experiment, compare the ***LD_50_*** value with the pesticide intake per bee during the same period, and examine whether it is possible to estimate and evaluate the pesticide administration-concentration in long-term field experiments from the ***LD_50_*** value. We will also evaluate the appropriateness of pesticide administration-concentrations in our long-term field experiments so far.

## Materials and Methods

### Field experimental method

We have conducted six long-term field experiments shown in Supplementary Table S2 to investigate the effects of neonicotinoid pesticides on bee (*Apis mellifera*) colonies (T. Yamada, et al., 2012, 2018_a_, 2018_b_, 2018_c_, 2018_d_; T. Yamada & K. Yamada, 2020; Yamada, 2020). In these six long-term field experiments, sugar syrup (SS) and pollen paste (PP), which are foods of honey bees, were used as a vehicle to administer pesticides, which were dinotefuran (DF), clothianidin (CN), fenitrothion (FT) and malathion (MT), to a honey bee colony. Supplementary Methods S1 to S3 show the details of materials used in long-term field experiments, the method of long-term field experiments, the method for counting the numbers of adult bees and capped brood, and the method for calculating the intakes of foods and pesticides such as the average daily intake of pesticide and food per bee, respectively.

### Calculation methods for foods and various pesticide intakes

#### 【Average intake of pesticide per day per bee】

The average intake of pesticide per day per bee, ***P_BD_***, is calculated by Equation (1), where values (***P_T_***, ***AB_T_***, ***I_T_***) other than ***P_B_*** in Equation (1) are reported in papers already published by T. Yamada, et al. (2018_a_, 2018_b_) and Yamada (2020). For details, see Supplementary Method S3.

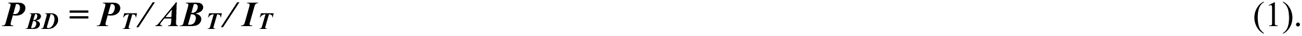

Where ***<u>P</u>_BD_*** is the average daily intake of pesticide per bee, ***P_T_*** is the total intake of pesticide during pesticide-administration period per colony, which is equal to the pesticide consumption during the period measured in the experiment, ***AB_T_*** is the total number of bees during pesticide-administration period per colony, which is calculated from the initial number of adult bees and the total number of new adult bees having eclosed from capped brood, and ***I_T_*** is the total number of days during pesticide-administration period to the colony, which is the total number of days from the start of pesticide administration to the stop of the administration. Here, ***P_T_*** and ***I_T_*** are directly obtained from the experiment, but ***AB_T_*** cannot directly.

#### 【Total number of bees during pesticide-administration】

Total number of bees during pesticide-administration period per colony, ***AB_T_***, can be obtained from Equation (2) under the assumption that the number distribution of capped brood is the same between aged 1 day to 12 days at the ***k^th^*** measurement date from the starting date of experiment and the brood have already ingested the pesticide at the end of experiment. For mor details, see T. Yamada et al., (2018_a_, 2018_b_).

The procedures to estimate the total number of adult bees during a certain period are as follows: Supposing that the number of adult bees at the first measurement (start of pesticide administration) is ***AB_0_***, the number of capped brood whose distribution aged 1 to 12 days is the same at the ***k^th^***measurement is ***CB_k_***, the number of days from the ***k^th^*** measurement to the (***k+1)^th^*** measurement is ***D_k_*** and the number of total observations is ***N***, ***AB_T_*** can be roughly obtained from the following equation: Where, it is assumed that the capped brood (***CB_N_***) present at the end of the administration period has already ingested food and pesticides.

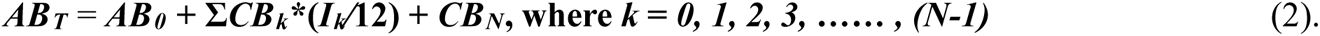

Where ***N*** is the total measurement number of times from the starting date of experiment (***N*** = 0) to the final date of experiment (***N*** = ***N***), ***k^th^*** is the measurement number of times from the starting date of experiment, ***AB_0_*** = the initial number of adult bees at the starting date of experiment, ***CB_k_*** is the number of capped brood at the ***k^th^*** measurement date, ***I_k_*** is the number of days of the interval between the ***k^th^*** measurement date and the (***k+1)^th^***measurement date. ***CB_N_*** is the number of capped brood at the final date of experiment when the brood are estimated to have already ingested the pesticide by then.

## Results

In six long-term field experiments conducted by T. Yamada et al. (2012, 2018_a_, 2018_b_, 2018_c_, 2018_d_) and Yamada (2020) as shown in Supplementary Table S2, pesticides were administered to bee colony via food (SS, PP). In only 2010-Experiment (T. Yamada et al., 2012), either dinotefuran or clothianidin was administered to honey bee colonies via both SS and PP. In other five long-term field experiments (2011/2012-Experiment reported in T. Yamada et al., 2018_a_, 2012/2013-Experiment reported in T. Yamada et al., 2018_b_, 2013/2014-Experiment reported in T. Yamada et al., 2018_c_, 2014/2015-Experiment reported in T. Yamada et al., 2018_d_, 2018-Experiment reported in Yamada, 2020) than 2010-Experiment, one kind of pesticides was administered to a honey bee colony via either SS or PP.

We excluded our first long-term field experiment, the 2010-Experiment, from this work for the following reasons: The first reason is that at the time of this experiment, there was no technology for measuring the number of adult bees and capped broods with high accuracy, and it was expected that the accuracy would be lower than that of the other five field experiments.

The second reason is that two types of pesticide-containing food (SS and PP) are administered at the same time, and the difference in intakes between SS and PP cannot be analyzed. Therefore, we analyzed using experimental data obtained from five long-term field experiments except for the 2010-experiment.

In this paper were assessed various intakes obtained by dividing the total intake of pesticides, SS and PP in honey bee colonies by the total number of adult bees during the administration period and/or the number of days of administration period for these five long-term field experiments. During the administration period to a honey bee colony in five long-term field experiments, Table 1 shows the experimental conditions of five long-term field experiments, and during each pesticide-administration period, the total number of adult bees in one colony and the total intakes of SS, PP and pesticide per colony, the total intakes per bee and the average daily intakes per bee. Table 1 was created by scrutinizing the original data of long-term field experiments, recalculating them, and making some corrections. Table 1 lists the experimental conditions and various intakes of foods (SS, PP) and pesticides during the pesticide administration period to a bee-colony. These intakes were obtained according to Supplementary Method 3.

**Table 1.**
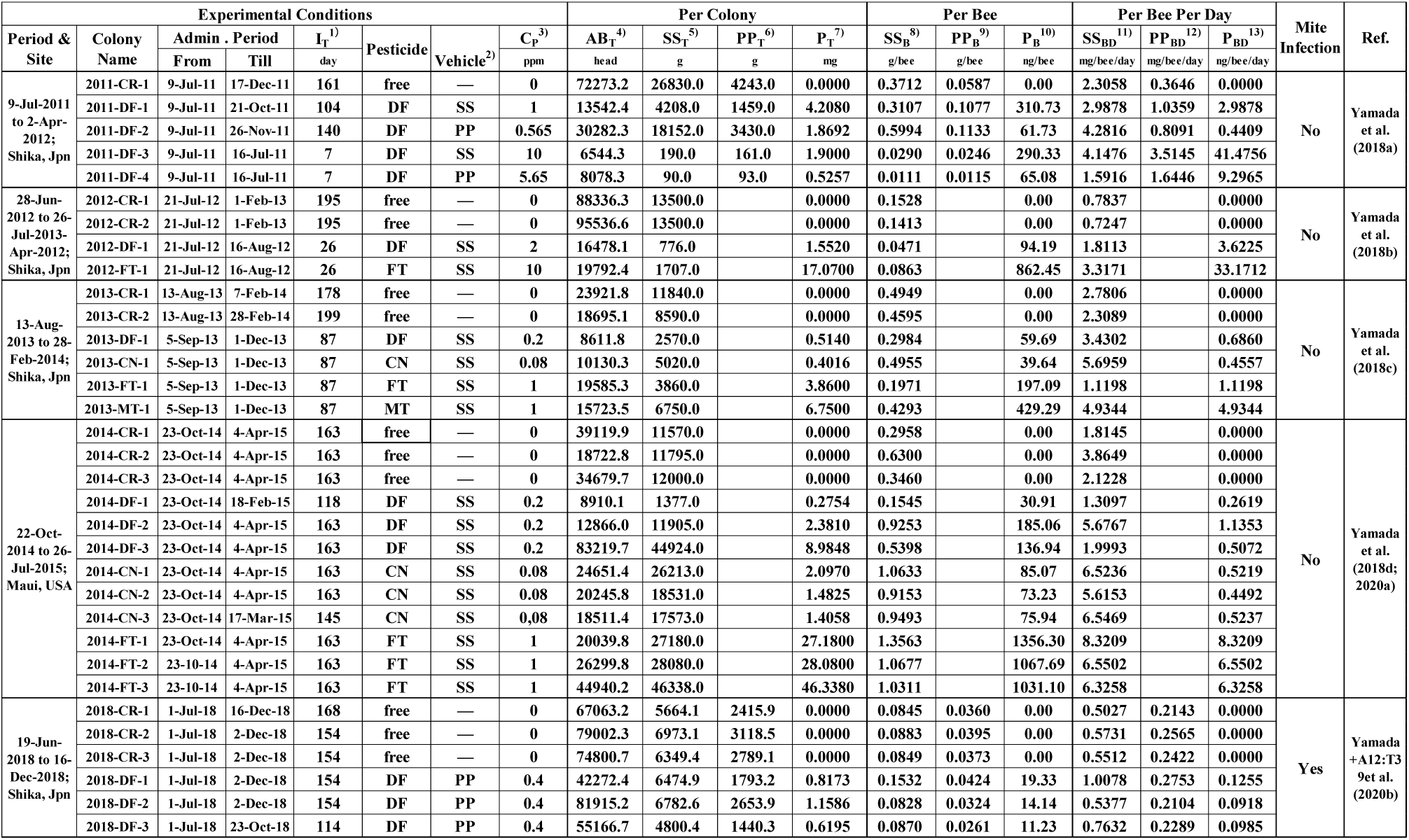
Intakes of sugar syrup, pollen paste and pesticide during the administration period to a bee colony in five long-term field experiments. Note (1): Each intake is recalculated under the assumption that each total amount of honey, pollen and pesticide is taken by bees during the period of its administration. Note (2): Since the figures in the table have been newly recalculated, they may differ slightly from the published figures. If there is no published data, leave it blank. Note (3): “free” = pesticide free, “DF” = dinotefuran, “CN” = clothianidin, “FT” = fenitrothion, “MT” = malathion. 1) ***I_T_*** denotes the number of days of total experimental intervals during pesticide administration. 2) “Vehicle” means food for administering the pesticide to colonies, SS and PP indicate sugar syrup and pollen paste, respectively. 3) ***C_P_*** denotes the concentration of the pesticide administered to the colony. 4) ***AB_T_*** denotes the total number of adult bees that would have ingested pesticides in the colony during pesticide administration to the bee colony (***I_T_***). 5) ***SS_T_*** denotes the total intake of sugar syrup ingested by the colony during pesticide administration to the bee colony. 6) ***PP_T_*** denotes the total intake of pollen paste ingested by the colony during pesticide administration to the bee colony. 7) ***P_T_*** denotes the total intake of the pesticide ingested by the colony during pesticide administration to the bee colony. 8) ***SS_B_*** denotes denotes the total intake of sugar syrup ingested by one bee during pesticide administration to the bee colony, ***SS_B_*** = ***SS_T_***/***AB_T_***. 9) ***PP_B_*** denotes the total intake of pollen paste ingested by one bee during pesticide administration to the bee colony, ***PP_B_*** = ***PP_T_***/***AB_T_***. 10) ***P_B_*** denotes the total intake of the pesticide ingested by one bee during pesticide administration to the bee colony, ***P_B_*** = ***P_T_***/***AB_T_***. 11) ***SS_BD_*** denotes the average daily intake of sugar syrup ingested by one bee during pesticide administration to the bee colony, ***SS_BD_*** = ***SS_T_***/***AB_T_***/***I_T_***. 12) ***PP_BD_*** denotes the average daily intake of pollen paste ingested by one bee during pesticide administration to the bee colony, ***PP_BD_*** = ***PP_T_***/***AB_T_***/***I_T_***. 13) ***P_BD_*** denotes the average daily intake of the pesticide ingested by one bee during pesticide administration to the bee colony, ***P_BD_*** = ***P_T_***/***AB_T_***/***I_T_***.

### Overall mean value in each targeted data group of the intakes of foods (SS, PP) and pesticides

Supplementary Table S5. lists each overall mean value of various average daily intakes for SS (***SS_BD_***), pollen paste (***PP_BD_***) and pesticides (***P_BD_***) ingested by one bee during the pesticide-administration period, and that of various total intakes for SS (***SS_B_***), pollen paste (***PP_B_***) and pesticides (***P_B_***) ingested by one bee during the period.

In calculating the overall mean value, the groups of all data, pesticide-free data, all pesticide data, dinotefuran (DF) administration data (both SS and PP; SS only; PP only), clothianidin (CN) administration data, fenitrothion (FT) administration data and malathion (MT) are used as the target data populations. Only SS was used as a vehicle for administering a pesticide to all but DF colony. From these overall mean values. In addition, the colonies where the neonicotinoid (DF, CN) was administered became extinct in every field experiment conducted by the authors (T. Yamada et al., 2012, 2018_a_, 2018_b_, 2018_c_, 2018_d_; Yamada, 2010), but the other colonies where the organophosphate (FT, MT) or no pesticide was administered did not always become extinct.

The overall mean values are as follows: For the total intakes of pesticides per bee during pesticide-administration, 0.42 g/bee of ***SS_B_*** and 0.048 g/bee of ***PP_B_*** for the group of all data (***SS_B-ALL_***, ***PP_B-ALL_***); 0.29 g/bee of ***SS_B_*** and 0.043 g/bee of ***PP_B_*** for the group of pesticide-free data (***SS_B-FR_***, ***PP_B-FR_***); 0.33 g/bee of ***SS_B_***, 0.066 mg/bee of ***PP_B_*** and 158 ng/bee of ***P_B_*** for the group of data where SS only contains DF (***SS_B-DF-SS_***, ***PP_B-DF-SS_***, ***P_B-DF-SS_***); 0.86 g/bee of ***SS_B_*** and 69 ng/bee of ***P_B_*** for the group of data where SS only contains CN (***SS_B-CN-SS_***, ***P_B-CN-SS_***); 0.75 g/bee of ***SS_B_*** and 903 ng/bee of ***P_B_*** for the group of data where SS only contains FT (***SS_B-FT-SS_***, ***P_B-FT-SS_***); 0.43 g/bee of ***SS_B_*** and 429 ng/bee of ***P_B_*** for the group of data where SS only contains MT (***SS_B-MT-SS_***, ***P_B-MT-SS_***).

For the average daily intakes of pesticides per bee during pesticide-administration, 3.1 mg/bee/day of ***SS_BD_*** and 0.8 mg/bee/day of ***PP_BD_*** for the group of all data (***SS_BD-ALL_***, ***PP_BD-ALL_***); 1.7 mg/bee/day of ***SS_BD_*** and 0.27 mg/bee/day of ***PP_BD_*** for the group of pesticide-free data (***SS_BD-FR_***, ***PP_BD-FR_***); 2.5 mg/bee/day of *SS_BD_*, 1.1 mg/bee/day of ***PP_BD_*** and 5.1 ng/bee/day of ***P_BD_*** for the group of all data where a vehicle (SS, PP) contains DF (***SS_BD-DF-ALL_***, ***PP_BD-DF-ALL_***); 3.1 mg/bee/day of ***SS_BD_***, 2.3 mg/bee/day of ***PP_BD_*** and 7.2 ng/bee/day of ***P_BD_*** for the group of data where SS only contains DF (***SS_BD-DF-SS_***, ***PP_BD-DF-SS_***, ***P_BD-DF-SS_***); 1.6 mg/bee/day of ***SS_BD_***, 0.6 mg/bee/day of ***PP_BD_*** and 2.0 ng/bee/day of ***P_BD_*** for the group of data where PP only contains DF (***SS_BD-DF-PP_***, ***PP_BD-DF-PP_***, ***P_BD-DF-PP_***); 6.1 mg/bee/day of ***SS_BD_*** and 0.49 ng/bee/day of ***P_BD_*** for the group of data where SS only contains CN (***SS_BD-CN-SS_***, ***P_BD-CN-SS_***); 5.4 mg/bee/day of ***SS_BD_*** and 11.3 ng/bee/day of ***P_BD_*** for the group of data where SS only contains FT (***SS_BD-FT-SS_***, ***P_BD-FT-SS_***); 4.9 mg/bee/day of ***SS_BD_*** and 4.9 ng/bee/day of ***P_BD_*** for the group of data where SS only contains MT (***SS_BD-MT-SS_***, ***P_BD-MT-SS_***). For the other intakes, see Supplementary Table S5.

From the above (Supplementary Table S5), the followings can be seen: The intake of SS is much higher than that of PP (e.g., ***SS_B-ALL_***/***PP_B-ALL_***; ***SS_B-FR_***, ***PP_B-FR_***; ***SS_BD-ALL_***/***PP_BD-ALL_***; ***SS_BD-FR_***/***PP_BD-FR_***). The intake of SS for CN is much higher than that for DF (***SS_B-DF-SS_***/ ***SS_B-DFCN-SS_***; ***SS_BD-DF-SS_***/ ***SS_BD-DFCN-SS_***). The total intake of the neonicotinoid (DF, CN) per bee is much higher than the organophosphate (FT, MT) (***P_B-DF-SS_*** & ***P_B-CN-SS_***/***P_B-FT-SS_*** & ***P_B-MT-SS_***), but for the average daily intake, the difference in the intake is not so high (***P_BD-DF-SS_*** & ***P_BD-CN-SS_***/***P_BD-FT-SS_*** & ***P_BD-MT-SS_***). The intake of SS for the pesticide-free colony is much lower than that for the pesticide-administration colony (***SS_B-FR_*** & ***SS_BD-FR_***/***SS_B-DF-SS_***, ***SS_B-CN-SS_***, ***SS_BD-DF-SS_*** & ***SS_BD-CN-SS_***), that of PP also is similar.

Let’s take ***SS_X-Y-Z_*** as an example to explain the subscripts. This is the intake of SS and the subscript is its explanation. The “X” in X-Y-Z of subscript means “per bee” for “B” and “per bee per day” for “BD”, and the “Y” denotes the kind of pesticide, viz., means “pesticide-free” for “FR” and “dinotefuran” for “DF”, etc., then, the “Z” denotes the kind of vehicles, means “sugar syrup” for SS, etc.

### Average daily intake of pesticide ingested by one bee, PBD

Figure 1 is a logarithmic plot of the relationship between the concentration of pesticide administered to the honey bee colony (***C_P_***) [ng (pesticide) ⁄ g(pesticide-containing vehicle)] and the average intake of pesticide per bee per day (***P_BD_***) [ng/bee/day], with ***C_P_*** on the horizontal axis and ***P_BD_*** on the vertical axis, for all data reported in previous our works on five long-term-field experiments. From Figure 1, it can be seen that ***C_P_*** and *P_BD_* is roughly expressed by the following equation.

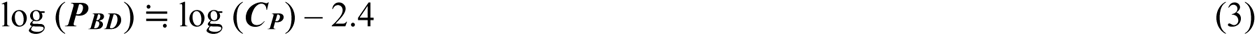

**Figure 1.**
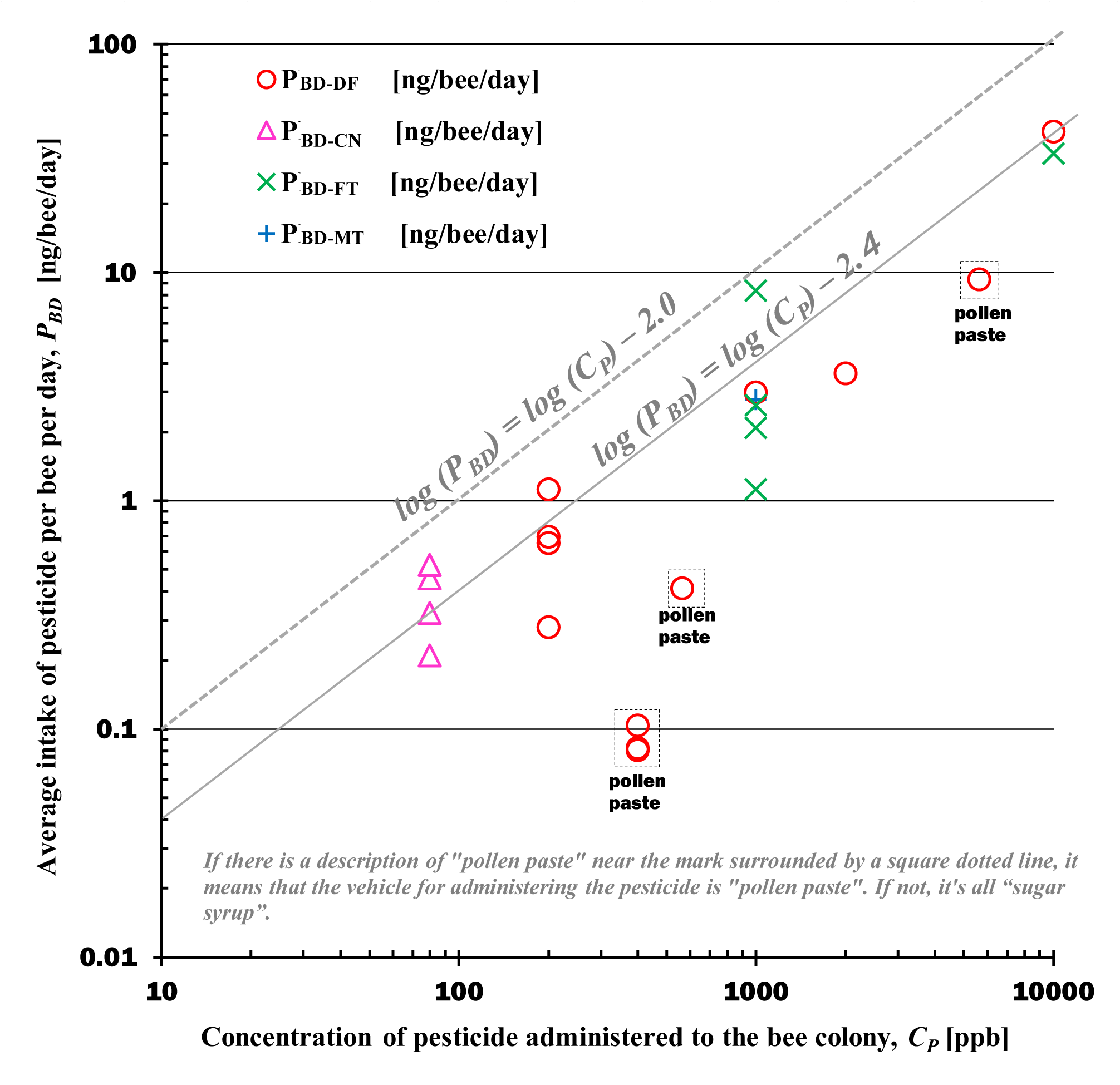
Relationship between the concentration of pesticide administered to the bee colony (***C_P_***) and the average daily intake of pesticide ingested by one bee (***P_BD_***). DF: Dinotefuran, CN: Clothianidin, FT: Fenitrothion, MT: Malathion. ***P_BD_*:** Average daily intake of a pesticide ingested by one bee during the administration period. ***PBD-DF***: Average daily intake of DF ingested by one bee during the administration period (***P_BD-DF_***). ***PBD-CN***: Average daily intake of CN ingested by one bee during the administration period (***P_BD-CN_***). ***PBD-FT*:** Average daily intake of FT ingested by one bee during the administration period (***P_BD-FT_***). ***PBD-MT*:** Average daily intake of dinotefuran ingested by one bee during the administration period (***P_BD-MT_***). The data surrounded by a dotted line in the upper rectangle labeled pollen paste in the figure shows the daily intake of pesticides by one bee when “pollen paste” containing pesticides is administered to a bee colony as a vehicle. All other data show the daily intake of pesticides per bee when pesticides are administered using sugar syrup as the vehicle.

That is,

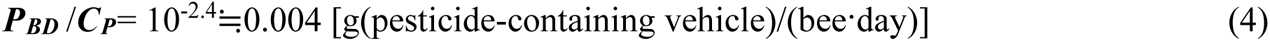

Parallel to equation (3), the logarithmic line that takes the maximum data of ***P_BD_*** is as follows.

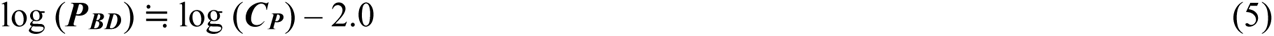

That is,

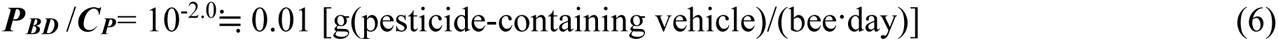

The followings can be foud from Table 1, Figure 1 and Equations (3) to (6): Broadly speaking, regardless of the type of pesticide administered, the higher the concentration administered, the higher the average pesticide intake per bee per day.

*P_BD_* administered with pesticides via pollen paste tends to be lower than *P_BD_* administered with pesticides via SS.

The relationship between ***P_BD_*** and ***C_P_*** is roughly expressed by Equation (3), regardless of the type of pesticide administered and the time and place of the experiment. It can be seen that ***P_BD_*** is less than or equal to Equation (5).

It can be seen from Equation (4) that the average pesticide intake (***P_BD_***) ingested by one bee per day is approximately 1/250 of the pesticide administration-concentration (***C_P_***) to the honey bee colony, and from Equation (6) that ***P_BD_*** is at most 1/100 of ***C_P_***. This suggests that environmental differences between open field experiments and closed laboratory experiments must be taken into account when determining the concentration of pesticides administered to honey bee colonies in long-term field experiments of honey bee colonies.

In addition, depending on the difference in vehicle, the pesticide intake per bee per day tends to be different. When a vehicle is “pollen paste (PP)”, the average intake of pesticide per bee per day is less than when “sugar syrup (SS)”.

### Total intakes of pesticide ingested by one bee during pesticide administration, PB

From Supplementary Figure S1 and Table 1, it can be said that the total intake of pesticide per bee (***P_B_***) appears to increase slightly with the increase in the concentration (***C_P_***) of pesticide administered to the honey bee colony, but broadly speaking, it can be almost independent of ***C_P_*** and, very roughly, constant. The grand average of all

Here, the average value of ***P_B_*** for each kind of pesticides and vehicles to administer the pesticide in five field experiments is as follows: The average of ***P_B_*** for dinotefuran via sugar syrup (***P_B-DF-SS_***) is 158.3 ng/bee; that for dinotefuran via PP (***P_B-DF-PP_***) is 34.3 ng/bee; that for clothianidin via sugar syrup (***P_B-CN-SS_***) is 68.5 ng/bee; that for fenitrothion via sugar syrup (***P_B-FT-SS_***) is 902.9 ng/bee; that for malathion (MT) via sugar syrup (SS) is 429.3 ng/bee. From the above results, it can be seen that the value of ***P_B-DF-PP_*** is about five times as much as the value of ***P_B-DF-SS_***, the value of ***P_B-DF-SS_*** is about three times as much as the value of ***P_B-CN-SS_***, and the value of ***P_B_*** of the neonicotinoid (DF, CN) for sugar syrup is extremely greater than that of the organophosphate (FT, MT).

### Median lethal dose of the bee (LD_50_)

The literature values of the median lethal dose of the bee (***LD_50_***) are shown in Supplementary Table S3, and the literatures cited in that table are listed in Supplementary Table S3_a_.

In Supplementary Table S3, various averages were calculated always from the original data, not from each average.

As the ***LD_50_*** values for comparing with the intake of pesticide, the overall mean value of ***LD_50_*** for acute oral toxicity for test durations of 24, 48 and 72 hours was adopted As the LD50 value for comparison with pesticide intake, the overall average value of acute oral toxicity LD50 over the 24-, 48-, and 72-hour test periods was adopted to reduce variability in the data. The oral toxicity data of ***LD_50_*** were used because the comparison with ***LD_50_*** was the intake of pesticides ingested by mouth obtained in long-term field experiments.

The overall mean values of acute oral toxicity LD50 at 24, 48, and 72 hours (mean test duration = 48 hours) were 28.2 ng/bee for dinotefuran, 54.0 ng/bee for clothianidin, 203.3 g/bee for fenitrothion, and 385.0 ng/bee for malathion.

### Ratio of P_BD_ to C_P_

In order to investigate the trend and extent of the effect of pesticide administration-concentration on the average daily intake of pesticide ingested by a single bee, the ratio of ***P_BD_*** to ***C_P_*** is plotted on a figure with the horizontal axis as ***C_P_***.

Supplementary Figure S2 shows the ratio of the average daily intake of pesticide per bee (***P_BD_***) during the pesticide administration period to the concentration of pesticide administered to the honey bee colony (***C_P_***). From Supplementary Figure S2, it can be seen that the ***P_BD_***/***C_P_*** value is almost constant, with the variations in the data, regardless of the pesticide administration-concentration and the type of pesticide. The arithmetic average value of all ***P_BD_***/***C_P_*** values is 3.5 mg/bee/day (horizontal dotted line in the figure). In addition, ***P_BD_***/***C_P_*** is almost equivalent to the intake of SS per bee per day from dimensional analysis. Therefore, comparing the arithmetic average value of all ***P_BD_***/***C_P_*** values and the average daily intake of SS per bee obtained from all data (see Supplementary Table S5), the arithmetic average value (3.5 mg/bee/day) of all ***P_BD_***/***C_P_*** values is found to be approximate to the average daily intake of SS per bee (3.1 mg/bee/day) (***SS_BD-ALL_*** in previous section, Supplementary Table 5).

### Total intake of food (SS, PP) per bee during the administration period

Supplementary Figure S3 shows the relationship between the administration-concentration of pesticide to the honey bee colony, ***C_P_***, and the total intake of food (SS, PP) per bee during the administration period, ***SS_B_*** & ***PP_B_***. In the field experiments, SS and PP are fed to a honey bee colony as a food or as a vehicle to administer a pesticide to a honey bee colony.

Here, we investigate how much food (SS, PP) one bee ingests during the administration period, whether the bee’s food intake is affected by the concentration of pesticides administered, and in particular, whether there is a difference between the intake of pesticide-free food and the intake of pesticide-containing food. Here, since the case where the pesticide concentration is 0 (zero) cannot be plotted on the logarithmic graph, in order to plot the pesticide-free data on the logarithmic graph, the pesticide-free concentration (***C_P_***) is assumed to be equivalent to 0.01 ppb.

The following can be seen from Supplementary Figure S3 and Table 1. Despite variations in the data, there is little difference between the total intake per one bee of pesticide-containing food and that of pesticide-free food. It should be noted that the total intake of PP ingested by one bee during the administration period is less than that of sugar syrup.

Food (SS or PP) intake by one bee appears to be almost constant regardless of the concentration of a pesticide administered to a honey bee colony (***C_P_***), although there are variations. That is, the overall mean value of all total intakes of SS and PP ingested by one bee during the administration period can be approximately 0.42 g/bee and 0.048 g/bee (Supplementary Table S5), respectively. And, the maximum total intakes of SS and PP ingested by one bee during the administration period can be approximately 1.4 g/bee and 0.11 g/bee (Table 1), respectively.

### Average daily intake of food (sugar syrup, pollen paste) per bee

Figure 3 shows the relationship between the administration-concentration of pesticide to the honey bee colony and the average daily intake of food (SS, PP) per bee during the administration period. The following can be seen from Figure 3 and Table 1.

**Figure 2.**
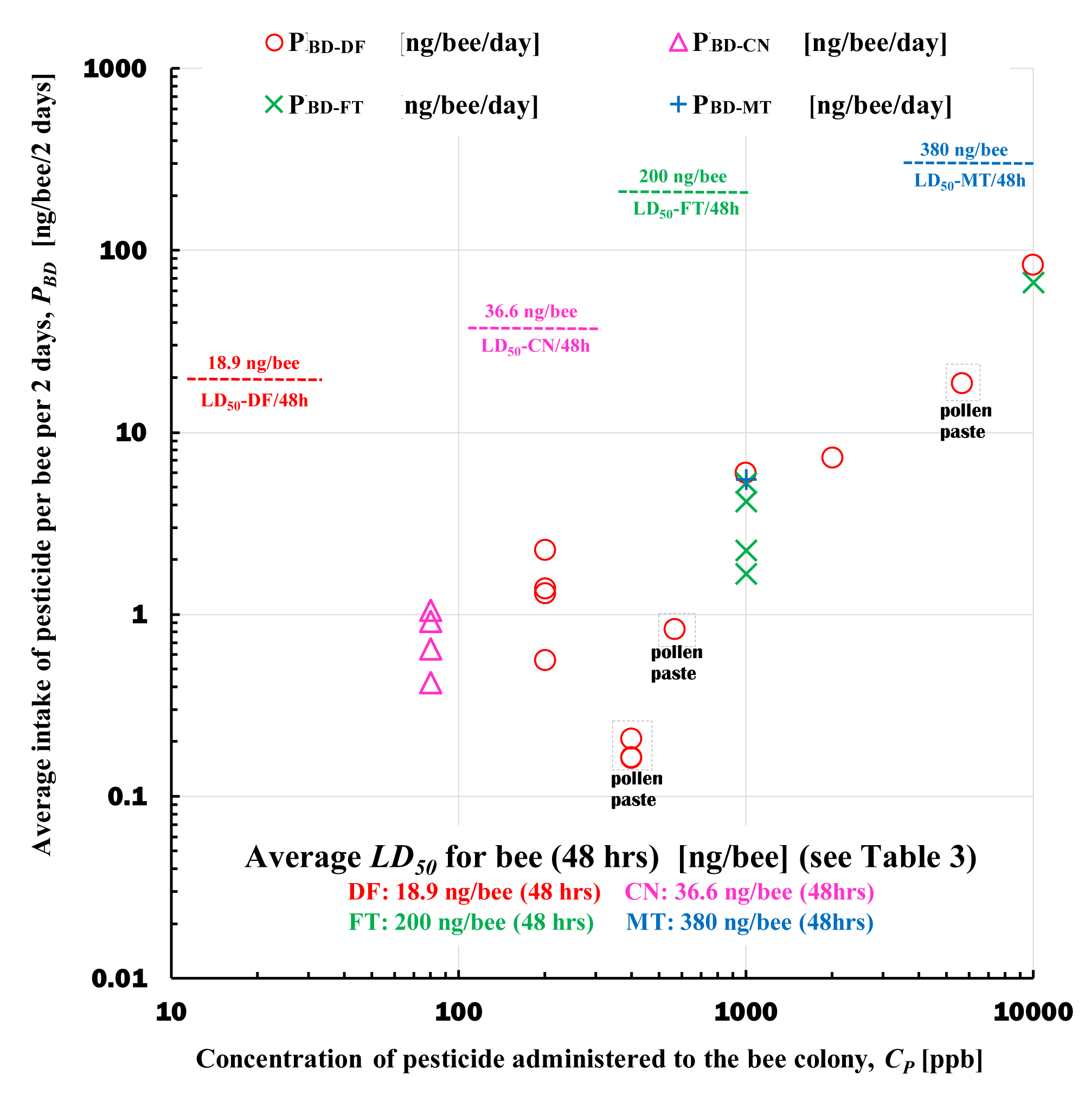
Comparison between average intake of pesticide per bee per two days (***P_BD_***) and median lethal dose of the bee (***LD_50_***). Dotted lines are the ***LD_50_*** values. “LD50-DF/48h”, “LD50-CN/48h”, “LD50-FT/48h” and “LD50-MT/48h” denote the ***LD_50_*** values of dinotefuran, clothianidin, fenitrothion and malathion for honeybee with measurement time 48 hours, respectively.

**Figure 3.**
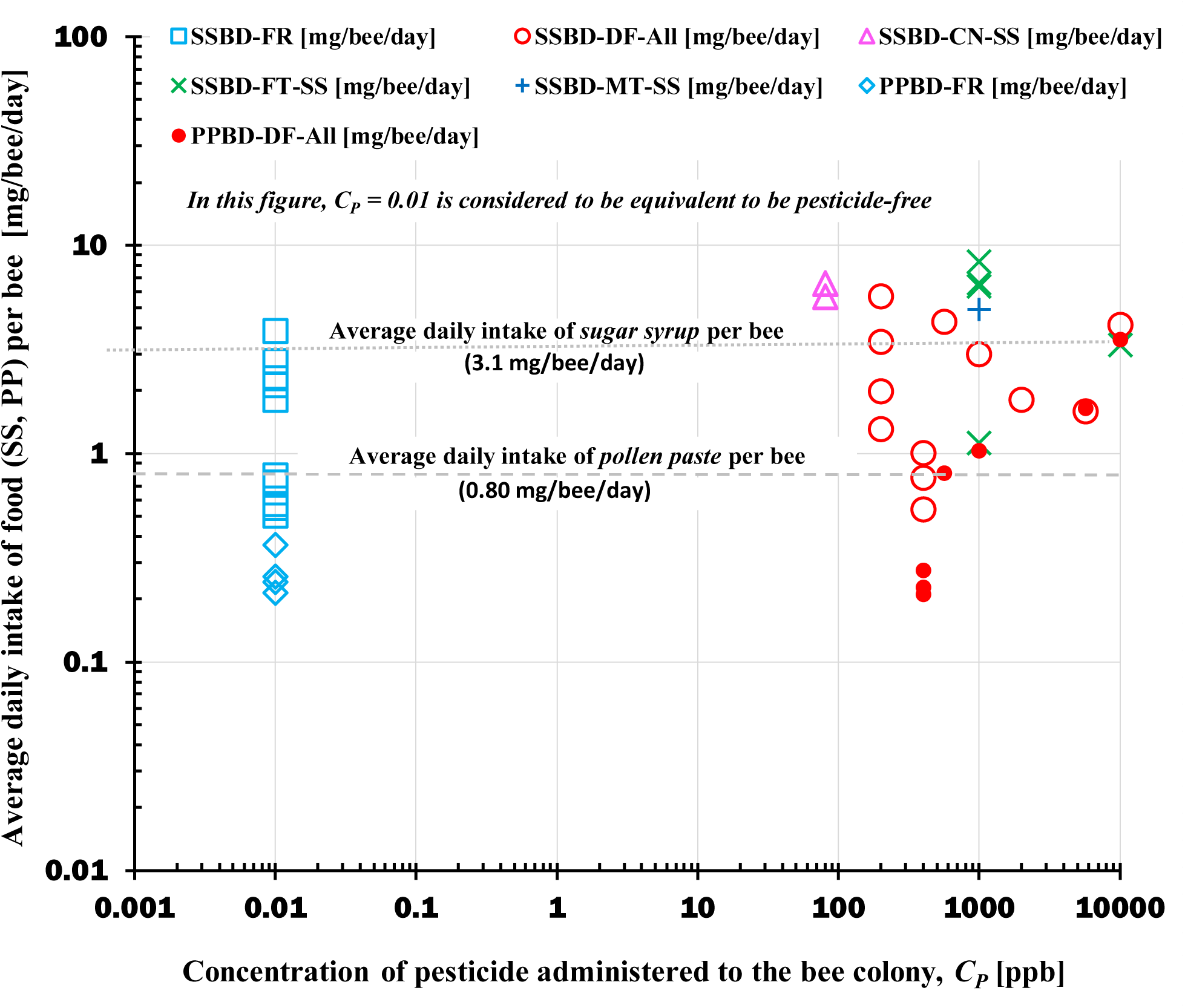
Relationship between the administration-concentration of pesticide to the bee colony and the average daily intake of food (sugar syrup, pollen paste) per bee during the administration period. SS: Sugar syrup, PP: Pollen paste, DF: Dinotefuran; CN: Clothianidin; FT: Fenitrothion; MT: Malathion. **SSBD-FR:** Average daily intake of pesticide-free SS per bee (***SS_BD-FR_***). **SSBD-DF-All:** Average daily intake of SS per bee when either SS or PP contains DF (***SS_BD-DF-All_***). **SSBD-CN-SS:** Average daily intake of SS containing CN per bee (***SS_BD-CN-SS_***). **SSBD-FT-SS:** Average daily intake of SS containing FT per bee (***SS_BD-FT-SS_***). **SSBD-FT-SS:** Average daily intake of SS containing malathion per bee (***SS_BD-FT-SS_***) **PPBD-FR:** Average daily intake of pesticide-free PP per bee (***PP_BD-FR_***). **PP_BD-DF-All_:** Average daily intake of PP per bee when either SS or PP contains DF (***PP_BD-DF-All_***). Average intake of sugar syrup per bee per day = 3.116 mg/bee/day; Average intake of pollen paste per bee per day =0.7997 mg/bee/day.

Despite variations in the data, there is little difference between the average intake per one bee of pesticide-containing food and that of pesticide-free food. The average daily intake of PP ingested by one bee is less than that of sugar syrup. Daily food (SS or PP) intake by one bee appears to be almost constant, regardless of the concentration (***C_P_***) of a pesticide administered to a honey bee colony, although there are variations.

That is, the average daily intake of sugar syrup ingested by one bee is approximately 3.1 mg/bee/day and that of PP approximately 0.8 mg/bee/day.

The value of 3.1 mg/bee/day which is the average daily intakes of sugar syrup ingested by one bee is approximately equal to the arithmetic average value of all ***P_BD_***/***C_P_*** [mg(pesticide-containing vehicle)/bee/day] values (3.5 mg/bee/day) described in Supplementary Figure S2. In addition, the maximum daily intakes of SS and PP ingested by one bee can be approximately 8.3mg/bee/day and 4.3 mg/bee/day, respectively (Table 1).

The arithmetic average value, 3.1 mg/bee/day, of all ***P_BD_***/***C_P_*** suggests that the concentration of pesticide administration in long-term field experiments can be estimated, as discussed earlier.

### Relationship between reduced C_P_ corrected by insecticidal activity,**【**C_P_)**】**, and various intakes Each pesticide has a different insecticide activity

In a long-term field experiment, the administration-concentration of each pesticide is adjusted so that the insecticidal activity of stink bugs is the same, and administered.

For example, 0.2 ppm of DF, 0.08 ppm of CN, 1 ppm of FT, and 1 ppm of MT have been found to have almost the same insecticidal activity against stink bugs. Therefore, when the pesticide administration-concentration is plotted on a graph with the horizontal axis, there is a problem that the insecticidal activity is different for different pesticides at the same point on the horizontal axis.

In order to solve this problem, it is necessary to express pesticide administration-concentrations with different insecticidal activity in dimensionless reduced administration-concentrations such that they have the same insecticidal activity at the same point on the horizontal axis.

Therefore, the dosage concentrations of various pesticides in long-term field experiments conducted so far were divided by reference concentrations (0.2 ppm for DF, 0.08 ppm for CN, 1 ppm for FT, and 1 ppm for MT) for each concentration, and dimensionless reduced administration-concentrations, [***C_P_***], were introduced. That is, [***C_P-DF_***] = ***C_P-DF_***/0.2, [***C_P-CN_***] = ***C_P-CN_***/0.08, [***C_P-FT_***] = ***C_P-FT_***, [***C_P-MT_***] = ***C_P-MT_***, where [***C_P-DF_***], [***C_P-CN_***], [***C_P-FT_***] and [***C_P-MT_***] are the reduced [***C_P_***], which is dimensionless, for dinotefuran, clothianidin, fenitrothion and malathion, respectively, and then, ***C_P-DF_***, ***C_P-CN_***, ***C_P-FT_*** and ***C_P-MT_***, which are dimensional, are the original administration-concentrations of pesticide for dinotefuran, clothianidin, fenitrothion and malathion, respectively.

Here, we introduce the results of a graph replotting in which ***P_BD_***, ***SS_BD_***, ***PP_BD_*** and ***SS_T_*** are replotted with the reduced administration-concentration, [***C_P_***], as the horizontal axis.

### 【Relationship between [***C_P_***] and ***P_BD_***】

Supplementary Figure S4 shows a graph with a reduced pesticide administration-concentration, [***C_P_***], on the horizontal axis and the average daily intake of pesticide per bee per day, ***P_BD_***, plotted on the vertical axis. This diagram includes the intake data, which are composed of the pesticide-free colony, the neonicotinoid (DF, CN) administration colony, and the organophosphate (FT, MT) administration colony.

From this figure, it can be seen that the average daily intake of pesticide per bee per day, ***P_BD_***, is expressed by the following equation.

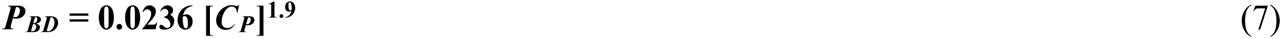

From this curve, the higher the reduced administration-concentration [***C_P_***], the higher the intake of pesticides. This suggests that these pesticides do not have a repellent effect on the ingestion of food containing pesticides by bees, but rather have a facilitating effect.

In addition, the average daily intake per bee ***P_BD_*** in the colony treated with the organophosphate deviated from this curve, indicating that the tendency was stronger as the reducing concentration [***C_P_***] increased.

In addition, the reduced ***C_P_*** of the colony administered the organophosphate deviates from this curve upward, and it can be seen that the tendency is stronger as the reduced concentration [*C*_P_] increases. That is, a colony that received the organophosphate at a concentration with the same insecticidal activity as the neonicotinoid survived even when it ingested a significantly higher amount of the organophosphate than the pesticide intake of the neonicotinoid-treated colony (when [CP] = 10). though all colonies receiving neonicotinoid are extinct.

From this fact, it can be inferred that the residual effect of the organophosphate was very short, and the insecticidal activity of the pesticide was extremely low when the colony administered the organophosphate ingested the organophosphate.

### 【Relationship between [***C_P_***] and ***SS_BD_***】

Supplementary Figure S5 shows a graph with a reduced pesticide administration-concentration, [***C_P_***], on the horizontal axis and the average daily intake of SS per bee, ***SS_BD_***, plotted on the vertical axis. From this figure, it can be seen that the intake per one bee ingested with the organophosphate (FT,。 In addition, at first glance in this figure, it appears that the average daily intake of SS per bee decreases as the reduced pesticide dose concentration [***C_P_***] increases.

However, if we look at this figure, excluding the data on the intake per one bee that ingested the organophosphate (FT, MT), it can be said that ***SS_BD_*** does not necessarily decrease as [***C_P_***] increases, and although there is a large variation, it is rather almost constant.

### 【Relationship between [***C_P_***] and ***PP_BD_***】

Supplementary Figure S6 shows a graph with a reduced pesticide administration-concentration, [***C_P_***], on the horizontal axis and the average daily intake of PP per bee, ***PP_BD_***, plotted on the vertical axis. It should be noted, however, that this figure does not include the intake per bee ingested with the organophosphate, but only the per bee intake in colonies that have not been treated with pesticides and colonies that have been treated with neonicotinoids (DF, CN).

### 【Relationship between [***C_P_***] and ***SS_T_***】

Supplementary Figure S7 shows a graph with a reduced pesticide administration-concentration, [***C_P_***], on the horizontal axis and the total intake of SS per colony, ***SS_T_***, plotted on the vertical axis.

Roughly speaking, this figure shows that the total intake of SS per colony, ***SS_T_***, decreases as [***C_P_***] increases. However, the intake per bee ingested with the organophosphate does not show an abnormal trend like PBD (Supplementary Figure 4) or SSBD (Supplementary Figure 5).

Broadly speaking, this figure shows that ***SS_T_***, which is the total intake of SS per colony, decreases with increasing [***C_P_***]. However, there is no abnormal trend in the intake per colony of organophosphate as in ***P_BD_*** (Supplementary Figure 4) and ***SS_BD_*** (Supplementary Figure 5), which are the average daily intake per bee.

## Discussion

### Guideline for setting appropriate pesticide concentrations

In order to investigate the effects of pesticides on individual bees, closed room experiments are suitable, which are easy to control the experimental environment, but in order to investigate the effects on these honey bee colonies in apiaries, open long-term field experiments, which are close to the actual apiary environment and extremely difficult to control the experimental environment, are suitable.

In field experiments, which are open environments, food that is often brought into the colony from outside other than the food (nectar, pollen) fed in the experiment (sugar syrup, pollen paste), and the differences in palatability (sugar syrup, flower nectar; pollen paste/bee bread) can be thought to be causing the difference in these intakes in field experiments.

In field experiments, which are open environments, food that is often brought into the colony from outside other than the food (nectar, pollen) fed in the experiment (sugar syrup, pollen paste), and the differences in palatability (sugar syrup, flower nectar; pollen paste/bee bread) can be thought to be causing the difference in these intakes in field experiments.

In pesticide administration experiments in field experiments, the administration-concentration of pesticide to bring the intake of pesticide-containing food by bees closer to the experimental target has been unknown until now. Therefore, without discussing the above differences from closed experiments in depth, it was often discussed whether the administration-concentration of pesticides was appropriate using the ***LD_50_*** value measured in laboratory experiments as an index in experiments to administer pesticides to bee colonies in the open field.

Here, we will try to propose a guideline for setting appropriate pesticide concentrations in field experiments.

When calculating and evaluating the appropriateness of pesticide administration-concentrations in long-term field experiments of honey bee colonies, it will be attempted to evaluate them at a median lethal dose (***LD_50_***) of bees under the assumption that the pesticide intake time of one bee in the field experiment is equivalent to the measurement time of ***LD_50_*** of honeybees.

In this paper, we compared the mean ***LD_50_*** value for bees of pesticides used in long-term field experiments with ***P_BD_*** values for various pesticide administration-concentrations in long-term field experiments with 48 hours when the test period was 48 hours. To assess the validity of estimating pesticide concentrations in long-term field experiments with ***LD_50_*** (test duration period: 48 hrs), we compared the ***P_BD_***×2 and ***LD_50_*** (48 hrs) intakes assuming one bee ingested the pesticide for two days (48 hrs) by doubling the *P_BD_* value.

Supplementary Table S3. shows the literature values of Median Lethal Dose (***LD_50_***) and various average values of these ***LD_50_*** data for Dinotefuran, Clothianidin, Fenitrothion and Malathion. The average ***LD_50_*** values of pesticides (48 hours) for acute oral toxicity and the ***P_BD_*** values obtained from long-term field experiments were used to evaluate the appropriateness of pesticide administration-concentrations for honey bee colonies in long-term field experiments.

Figure 2 shows the comparison between the average intakes of pesticide per bee per two days (48 hrs) and the average median lethal dose of the honeybee for acute oral toxicity at 48 hrs (AOT-LD50-48h in Supplementary Table S3). From Figure 2, when the concentration (***C_P_***) of pesticides administered to honey bee colonies in long-term field experiments is 5650 ppb in PP and 10000 ppb in sugar syrup, the average intake of pesticides per bee per two days (***P_BD_***x2) is equal to or greater than the ***LD_50_*** values.

The results were almost the same as above when compared with the overall average LD50 at 24, 48, and 72 hours (28.2 ng/bee for dinotefuran, 54.0 ng/bee for clothianidin, 203.3 g/bee for fenitrothion, and 385.0 ng/bee for malathion).

As described above, when the administration-concentration to bee colonies was high, bee colonies became extinct in a short period of time in previous field experiments (e.g. T. Yamada, et al., 2012). On the other hand, below these high concentrations, the ***P_BD_***x2 value is below the ***LD_50_*** value. This confirms the fact that in long-term field experiments, when pesticides were administered at concentrations lower than these high concentrations, honey bee colonies did not become extinct in a short period of time, but became extinct after showing the appearance of CCD, which is thought to be due to chronic toxicity (e.g. T. Yamada, et al., 2012).

These results suggest that Equations (3) (Figure 2) and (7) (Supplementary Figure S4) can be used as a guideline for setting appropriate pesticide concentrations in field pesticide administration experiments. In addition, if the insecticidal activity of each pesticide used is known and any one of DF, CN, FT and MT is included in it, it seems that the pesticide concentration can be set with higher accuracy by using Equation (7) than Equation (3).

### Simplified method to estimate the pesticide administration-concentration

The arithmetic average value, 3.5 mg/bee/day (Supplementary Figure S2), of all ***P_BD_***/***C_P_*** suggests that the concentration of pesticide administration in long-term field experiments can be easily estimated, as discussed earlier.

Now, assuming that the ***LD_50_*** value (24 h) is the maximum value of the target administration-concentration of pesticide, and if the maximum administration-concentration of pesticide is ***P_max_***, the ***P_BD_***/***C_P_*** is almost constant (average value 3.5 mg/bee/day) (Supplementary Figure S2), and the ***P_BD_***/***C_P_*** is equivalent to the volume (***V***) of the vehicle, so ***P_max_*** ×***V*** = ***LD_50-24h_*** holds.

Since ***P_BD_***/***C_P_*** = ***V*** = 3.5 mg/bee/day = 3.5×106 ng/bee/day, ***P_max_*** = ***LD_50-24h_/3.5*** [ppm], where it is given by ***LD_50-24h_*** [ng/bee]. That is, in this case, since the pesticide concentration of ***P_max_*** acts the same acute toxicity as ***LD_50_***, the pesticide administration-concentration can be set with reference to this value in a long-term field experiment.

In the past, long-term field experiments did not know the appropriate pesticide administration-concentration, and the pesticide administration-concentration was sometimes too high or too low, but the simplified estimation method described in this paper can provide a rough guideline for setting an appropriate pesticide concentration in long-term field experiments.

### Is the pesticide-containing food repellent or preferent for honey bee?

The overall mean value of the total intakes of SS ingested by one bee during the administration period in the field for the pesticide-free colonies (0.29 g/bee) are lower than that for all colonies (0.42 g/bee). The overall mean value of the average daily intakes of SS ingested by one bee during the administration period also is similar (1.67 mg/bee/day for pesticide-free colonies; 3.12 mg/bee/day for all colonies). Similarly, the overall mean values of PP for the pesticide-free colonies are lower than that for all colonies.

From the above results, the honey bee seems not to be repellent to the neonicotinoid (DF, CN) and the organophosphate (FT, MT), but seems to be rather even palatable to. In addition, the honey bee is most palatable to CN (0.86 g/bee for the total intake of SS; 6.10 mg/bee/day for the average daily intake of SS).

### Intakes of pesticide ingested by one bee

For honey bee colonies treated with neonics, the end of pesticide administration is the time of colony extinction, suggesting that the total pesticide intake (***P_B_***) of one bee until colony extinction is almost constant, independent of the pesticide administration concentration (***C_P_***) (Supplementary Figure S1). In other words, it is presumed that these intakes are almost the same as the total intake of pesticides until the honey bee colony becomes extinct when neonicotinoids are continuously ingested as chronic toxicity.

According to the overall mean values of the total intake per bee in Supplementary Table S5, it can be seen that the overall mean value of *P_B-DF-SS_* (158 ng/bee) is about 4.6 times as much as the value of *P_B-DF-PP_* (34 ng/bee), the that of *P_B-DF-SS_* is about 2.3 times as much as the value of *P_B-CN-SS_* (69 ng/bee), and that of the neonicotinoid (DF, CN) for sugar syrup is extremely lower than that of the organophosphate (FT, MT).

Judging from the fact that all the colonies having ingested the neonicotinoid (DF, CN) became extinct during pesticide administration but the colonies having ingested the organophosphate did not always (T. Yamada et al., 2012; 2018_a_; 2018_b_; 2018_c_; 2018_d_; Yamada, 2020), the intake of the neonicotinoid can be regarded as the intake till the colony extinction.

The difference of *P_B-DF-SS_* and *P_B-DF-PP_* can be due to that SS (honey) is the staple food of bees and that PP (pollen) is the staple food of larva and pupae (brood). As principal organs and functions are formed during the larval and pupal stages, the neonicotinoid which is long-lasting continues to affect adversely on the organs and functions, and as a result, PP with the neonicotinoid can destroy the bee colony at lower intake than SS. The difference of *P_B-DF-SS_* and *P_B-CN-SS_* is probably due to the difference in insecticidal activity, judging from the insecticidal activity of CN is about 2.5 times of that of DF.

According to the overall mean values of the average daily intake per bee in Supplementary Table S5, it can be found that the overall mean value of *P_BD-DF-SS_* (7.2 ng/bee/day) is about 3.6 times as much as the value of *P_BD-DF-PP_* (2.0 ng/bee)), the that of *P_BD-DF-SS_* is about 15 times as much as the value of *P_BD-CN-SS_* (0.49 ng/bee), The difference of *P_BD-DF-SS_* and *P_BD-DF-PP_* can be due to the same reasons in the total intakes of *P_B-DF-SS_* and *P_B-DF-PP_*. However, the difference of *P_B-DF-SS_* and *P_B-CN-SS_* cannot be confidently explained for the reason, although differences in insecticidal activity must have an effect.

## Conclusions

Depending on whether bees prefer SS and PP administered to a honey bee colony or on whether they prefer nectar and pollen collected from nature, the intake of SS and PP may differ in long-term field experiments. As a result of analysis based on experimental data in different environments, that is, in different areas (Shika Town in Japan, Maui Island in Hawaii) and at different times (experimental years), it is possible to clarify the relationship between the pesticide administration-concentration and bee intake in the open environment of long-term field experiments, although roughly, and provide a modest but somewhat meaningful guideline for determining pesticide administration-concentration in long-term field experiments. I believe it was given in this paper.

## Author Contributions

Toshiro Yamada conceived and analyzed the data, prepared the figures and/or tables, authored and reviewed drafts of the paper, and approved the final draft.

## Acknowledgments

Dr. Ken Hashimoto (Okayama, Japan) gave me encouragement.

## Supplementary Materials

**Supplementary Method S1.** Preparation and procedures for field experiments.

**Supplementary Method S2.** Counting methods of the numbers of adult bees and capped brood.

**Supplementary Method S3.** Calculation methods for the intakes of foods and pesticides.

**Supplementary Figure S1.** Relationship between the concentration of pesticide administered to the bee colony (***C_P_***) and the total intake of pesticide per bee (***P_B_***).

**Supplementary Figure S2.** Ratio of the average daily intake of pesticide per bee (*P_BD_*) during the pesticide administration period to the concentration of pesticide administered to the bee colony (*C_P_*).

**Supplementary Figure S3.** Relationship between the administration-concentration of pesticide to the bee colony and the total intake of food (sugar syrup, pollen paste) per bee during the administration period.

**Supplementary Figure S4.** Relationship between the reduced concentration of pesticide administered to the bee colony, [***C_P_***], and the average intake of pesticide per bee per day (***P_BD_***) during pesticide administration.

**Supplementary Figure S5.** Relationship between the reduced concentration of pesticide administered to the bee colony, [***C_P_***], and the average intake of sugar syrup per bee per day (***SS_BD_***) during pesticide administration.

**Supplementary Figure S6.** Relationship between the reduced concentration of pesticide administered to the bee colony, [***C_P_***], and the average intake of pollen paste per bee per day (***PP_BD_***) during pesticide administration.

**Supplementary Figure S7.** Relationship between the reduced concentration of pesticide administered to the bee colony, [***C_P_***], and the total intake of sugar syrup per colony (***SS_T_***) during pesticide administration.

**Supplementary Table S1.** Comparison between long-term field experiments and ***LD_50_*** measurement experiments in the investigation of the effects of pesticides on bees.

**Supplementary Table S2.** Experimental conditions of six long-term field experiments for honeybee colonies.

**Supplementary Table S3.** Literature values of median lethal doses (***LD_50_***) for the honeybee and various average values of the ***LD_50_*** data for dinotefuran, clothianidin, fenitrothion and malathion.

**Supplementary Table S3a.** List of literatures cited in Supplementary Table S3.

**Supplementary Table S4.** Example data for intakes of food and pesticide.

**Table S5.** Overall mean value in each data group of ***SS_B_***, ***PP_B_*** and ***P_B_*** and that in each data group of ***SS_BD_***, ***PP_BD_*** and ***P_BD_*** during the pesticide administration period.

## Supplementary Materials for

### This PDF file includes

Materials and Methods

Figures. S1 to S7

Tables S1 to S5

References

## 1. Supplementary Materials and Methods

### 1.1 Supplementary Method S1

(Preparation and procedures for field experiments)

#### 1.1.1 Materials

The following pesticides were used to examine their impacts on a bee colony.

##### 【Neonicotinoid pesticides (Neonics)】

1) **Dinotefuran (DF):** Starkle Mate® (10% DF; Mitsui Chemicals Aglo, Inc., Tokyo, Japan).
2) **Clothianidin (CN):** DANTOTSU® (16% clothianidin; Sumitomo Co. Ltd., Tokyo, Japan).

##### 【Organophosphate pesticides (OPs)】

1) **Fenitrothion (FT):** SUMITHION^®^ emulsion (50% fenitrothion; Sumitomo Co. Ltd., Japan).
2) **Malathion (MT):** MALATHON® emulsion (50% malathion; Sumitomo Co. Ltd., Japan). The following honey bee foods were used as a vehicle to administer the pesticides.

##### 【Vehicles to administer the pesticides】

1) **Sugar syrup (SS):** Granulated sugar was purchased from the Japan Beekeeping Association (http://www.beekeeping.or.jp/) and was composed of 99.7988% purified sugar (granulated sugar), Sodium chloride (salt) 0.1% or more, L-lysine hydrochloride 0.1% or more, and food dye (Blue No. 2) 0.0012% or more). A total of 20 kg of granulated sugar and 13.33 kg of hot water at about 75 ℃ were mixed in a 50 L plastic tank, and then 60% SS of granulated sugar was produced.
2) **Pollen paste (PP):** In total, 25 kg of pollen from Spain was purchased from Tawara Apiaries Co., Ltd., Kobe, Japan (https://tawara88.com/about.html). The pollen was used after it was lightly ground with “Kona Ace A-7” flour milling equipment manufactured by Kokkousha Co., Ltd., Nagoya, Japan (http://www.kokkousha.co.jp/). The viscosity of PP was determined by the ratio of SS in PP, the particle size of the pollen, and the temperature of the PP. Pollen pastes of various ratios of SS to PP were prepared and the ratio of SS to PP was determined that did not fall off when the tray filled with PP was turned over. As a result, 60% pollen by weight and 40% SS by weight was confirmed to be an appropriate ratio. Pesticide-free PP and PP containing pesticide of desired concentration were prepared in 10 kg each. PP (6 kg of pollen and 4 kg of SS) was prepared in a large plastic bucket by kneading with a drill screwdriver with stirring blades used for high viscosity at a low speed until achieving a uniform paste.

#### 1.1.2 Methods for preparing pesticide dosing vehicles

The concentration of each pesticide in a vehicle was adjusted so that it has the same insecticidal activity in the same field experiment. For examples, 0.2 ppm of DF, 0.08 ppm of CN, 1 ppm of fenitrothion and 1 ppm of malathion have the same insecticidal activity. In our field experiments, the concentration of pesticide was adjusted to match these ratios.

##### 【Preparation of SS with pesticide of a desired concentration】

It was produced using an SS solution containing the desired concentration of pesticide and 60% by weight of sugar. The prepared SS-containing pesticide was placed in a 10 L container, shaded from light with a black bag, and stored in the refrigerator.

##### 【Preparation of the tray filled with PP containing a desired pesticide concentration】

In order to adjust pesticides to desired concentration in PP which consists 60% pollen and 40%SS in weight, pesticide concentration in PP is adjusted only by SS containing the pesticide. The 300 g PP containing the desired pesticide concentration, after being weighed in an upper plate balance scale (accuracy ± 1 g), was packed into a foamed polystyrene tray. Then, the PP tray filled with pollen paste was wrapped by thin plastic film to prevent the evaporation of moisture from the tray and stored in a refrigerator to prevent alteration of the PP. The pesticide-free PP tray was stored in the refrigerator compartment, and the PP tray containing a pesticide was stored in the freezer compartment.

#### 1.1.3 Items to be measured in field experiments and their precautions

In long-term field experiments on honeybee colonies, in order to understand the health of the colony in the hive-box, it is necessary to determine the presence or absence of a queen bee, the change in the number of adult bees over time, the change in the number of eggs laid over time, the change in the number of larvae over time, the change in the number of pupae over time, the change in the number of dead bees over time, the change in the number of drone bees over time, the number of wax moth larvae and the hive beetle (rarely found in Japan). It is necessary to grasp the presence or absence of pests such as mites and parasites, and the presence or absence of pathogen such as foulbrood and chalk disease.

In addition, it is necessary to protect against attacks from outside the hive-box, such as hornets, bears and toads. By the way, when starting a long-term field experiment, it is natural to obtain a honey bee colony that does not suffer from foulbrood, chalk disease, etc., and does not have mites, and pay close attention to these diseases during the experiment. During field experiments, it is possible to obtain a healthy honey bee colony at the beginning of the experiment, and to select and maintain the experimental environment, so that it is possible to protect against the infection of these pathogens and attacks from pests and vermin to some extent.

Therefore, in order to know the changes in the state of the honey bee colony in long-term field experiments, it is important to know the presence or absence of a queen bee, the over-time change in the number of adult bees, that of eggs laid, that of larvae, that of pupae, and that of dead bees. By the way, the over-time change in the number of eggs laid and that in the number of larvae can be approximately represented by the change in the number of capped brood (pupae) that are easy to measure.

In addition, even if it has not occurred at the start, it is necessary to record abnormalities inside and outside the colony that occur suddenly during the experiment, although not regularly (presence or absence of beetles, mites, chalk disease, queen cells, etc.).

Therefore, in the long-term field experiments, the presence or absence of queen bees, the number of adult bees, the number of eggs laid, the number of capped brood (pupae), and the number of dead bees were measured at each discrete observation experiment.

And various unexpected events, such as attacks of beetles, ticks and wasps, the presence of queen cells, were recorded in as much detail as possible at the time of occurrence. For details, refer to the previous papers (Yamada et al., 2018_a_, 2018_b_, 2018_c_, 2018_d_; Yamada T. & Yamada K., 2020; Yamada, 2020).

Of these measurements, the over-time change in the number of adult bees and that of capped brood are the most important for grasping the state-changes of the bee colony. In particular, the longevity of a honey bee colony is a very important indicator for estimating the length of the colony’s survival.

Yamada Y. & Yamada T. (2019) have proposed a mathematical model that can estimate the apparent longevity of a honey bee colony, based on the number of adult bees and the number of capped brood, which can be measured experimentally. Naturally, the accuracy of estimating the apparent longevity is strongly dependent on the accuracy of the measurement of the numbers of adult bees and capped brood. In addition, both numbers of adult bees and capped brood are closely related to each other throughout the honeybee life cycle.

If the measurement accuracy of either number is poor, a contradiction will arise between these two measurements. For example, when some of adult bees have gone out foraging, the number of adult bees will underestimate the actual number existing in the honey bee colony. In this case, because there is an unreasonable difference in numbers between the two, apparent longevity cannot be accurately obtained when solving simultaneous equations in a mathematical model, or cannot be sometimes even determined because the simultaneous equations cannot be solved.

Therefore, experiments are conducted while paying attention to the following points to measure accurately the numbers: The experiment is started immediately after dawn to minimize the number of adult bees out of the hive-box. The comb-frame with bees is pulled out from the hive-box as gently as possible, when taking picture, to prevent the bees from flying away, and is gently set on the photo stand. The experiment is conducted while avoiding a windy or rainy day, judging from the weather forecast, because of preventing the bees from becoming restless. In more details, see the previous paper (Yamada, 2020).

#### 1.1.4 Preparations for field experiment

Field experiments were conducted six times as shown in Supplementary Table S2 which summarizes the experimental period, the experimental site, the research object, the circumstances around experimental site, the experimental conditions, the experimental methods and the published articles, five times in Midwest Japan (Shika-machi, Hakui-gun, Ishikawa prefecture, Japan) and one time in Maui in Hawaii, U.S.A. Of the five field experiments in Midwest Japan, four (Yamada et al., 2012, 2018_a_, 2018_b_, 2018_c_) were conducted at the same location, and the remaining one (Yamada, 2020) was conducted at a location near the four experiments.

Each hive-box was arranged at intervals of 80 cm, with its entrance facing south, on a stand of about 20 cm in height, and then a large PVC tray with many perforations of 3mm in diameter to prevent water from collecting was placed between the hive-box and the stand for the measurement of the number of dead bees.

While keeping in mind that the observation results of the photo images are analyzed, various measures were taken to prevent errors and errors in field experiments. For example, in order to be able to infer the experimental contents of the honey bee colony in the hive-box, a large display tag was attached at the top of the front of each hive. A label was attached to the bottom of the hive-box or on four walls, displaying which part of the hive-box it was. Moreover, several labels displayed so that the position in the hive-box of each comb-frame and the front and back of the comb-frame could be understood were attached to the pier of the comb-frame. Further, the name of the hive-box where sugar syrup (SS) and pollen paste (PP) would be administered was described on a container with SS and a tray with PP in order to prevent the administration mistakes.

#### 1.1.5 Field experimental procedures

The most important data in the field experiment of honeybee colony are both the numbers of adult bees and capped brood. To obtain these data as accurately as possible, the field experiment was carried out by the following procedure.

STEP-1) To avoid being stung by honey bees, take proper protective measures before starting the experiment.
STEP-2) Before the start of experiment, prepare several sets of apicultural tools, a laboratory notebook, observation data recording papers, containers with SS, trays with PP, camera, a few empty hive-boxes, several pairs of tweezers or the like.
STEP-3) Take a picture of the experiment site (entire hive-box) before the experiment (check whether abnormal conditions have occurred).
STEP-4) From a large tray placed under the hive-box, count the number of dead bees while picking them up with tweezers, and record the number of dead bees outside the hive-box on a record paper. Discard the ones you have counted from the tray. Furthermore, the number of dead bees in the hive-box is counted during the internal inspection of the hive-box and entered on a record sheet as the number of dead bees in the hive-box. The total number of dead bees inside and outside the hive-box is the number of dead bees in the target colony (hive).
STEP-5) Before inspecting the inside of the hive-box, take a picture of the front of the hive-box (record whether the target hive-box to be observed and measured is correct).
STEP-6) While opening the hive-box and gently returning the bees attached to the hive cloth, remove the fabric that covered the top of comb-frames. After that, quickly, take a picture of the entire upper part of comb-frames in the hive-box,
STEP-7) Remove the tray containing PP and the container containing SS (tray and feeding frame) that were placed in the hive-box during the previous observation experiment from the hive-box, and then measure the remaining amount of PP and that of SS with the accurate balance of the upper plate. When removing the container from the hive-box, gently return the bees attached to the container to the hive-box.
STEP-8) Gently remove the comb-frame with bees from the hive-box in the order of number (in order from left to right toward the front of the hive-box), take a picture of both sides of each comb-frame. When you are taking a picture, if you find a queen bee, after taking a picture of the queen bee, put it in the queen cage and once isolate it. When finishing the photograph of the comb-frame, put the comb-frame in the spare empty hive-box prepared beforehand, and the lid of the hive-box is closed. The queen bee in the queen cage is placed at the top of comb-frame in the spare hive-box.
STEP-9) After photographing both sides of the comb-frame with bees and placing all of them in the spare empty box, photograph the four sides and the bottom of the hive-box to determine the number of adult bees remaining in the hive-box. If you can’t find a queen bee when you’re taking a picture of a comb-frame, carefully check for the presence or absence of the queen bee, and if present, take a picture. If adult bees are outside the hive-box as well as inside the hive-box (sometimes near the entrance of the hive-box at tropical nights), take a picture to count the number. After taking pictures of all the adult bees, look for the queen bee in the original hive-box again when you cannot find it, and then transfer all the remaining bees from the original hive to the spare box and clean the original hive-box.
STEP-10) Even if the queen bee cannot be found, the search for the queen bee is completed after carefully checking again whether there is a queen bee on all comb-frames and all the walls (including the bottom) inside the hive-box. Then, do the next task, which is to take a photo of the capped brood.
STEP-11) Take out the comb-frame with bees that had been placed in the spare hive-box in the number order, and return all the bees attached to the comb-frame to the original hive-box, while shaking the honey bees from the comb-frame. After taking pictures of both sides of the comb-frame without honey bees in the order of the comb-frame number, the comb-frame is also returned to the original position of the original hive-box. After a few comb-frames are returned to the original hive-box, return and release the queen bee in the queen cage, which is placed at the top of comb-frame in the spare hive-box, into the original hive-box. The photos of comb-frame without honey bees are used to count the number of capped brood.
STEP-12) After taking pictures of all comb-frames without honey bees, return the remaining honey bees in the spare hive-box to the original hive-box.
STEP-13) Various anomalies such as queen cells, *Varroa* mites, and the wax-moth larvae (wax worms) are often encountered when photographing comb-frames without honey bees. When you find such unusual situations, record them in photos as evidence and at the same time describe them in detail in your laboratory notebook.
STEP-14) If the queen bee is not found by all means, check in the next experiment. If it is not found next time, by the presence or absence of queen cells and by observing in detail the situation of oviposition, etc., the presence or absence of a queen bee is determined.
STEP-15) After covering the cloth on the top of comb-frames, put new SS and PP in the target hive-box, the measurement is ended with the lid of the hive-box.
STEP-16) After completion of the experiment, the test contents are confirmed and described in the laboratory notebook.

### 1.2 Counting methods of the numbers of adult bees and capped brood

#### 1.2.1. Circumstances for developing the automatic counting software and its issues

Using the photographs taken as described above, it is possible to determine the number of adult bees, the number of capped brood, and the number of mite-damaged bees. Counting these numbers, however, from photographs is an extremely difficult task. Therefore, with the cooperation of Mr. Yoshiki Nagai of Nanosystem Co., Ltd., Kyoto, Japan (http://nanosystem.jp/firm.htm), which develops image processing software, the author has developed computer software in 2012 that automatically counts these numbers from photographic images, and improvements have been made to the software since. As a result, the accuracy and operation of the counting have been greatly improved. The counting time and counting accuracy using this software are significantly improved compared to the amount of work required to count by hand directly from the original photo.

However, these counts are still inaccurate because of honey bee overlap, out-of-focus images, significant differences in contrast in the photographic images, misjudgments of the counting targets, and so on. In this software, it is also possible to manually correct after automatic counting. Therefore, after automatic counting, the counting mistakes due to the automation are corrected by manual operation while zooming into the image, and the corrected number becomes the final data. Thus, errors still emerge in automation when using this software, but by using manual operation, it is possible to correct the counting errors. The improvements in the counting accuracy and speed under this software contributed greatly to the improvement of the data analysis accuracy and the speed in the field experiments. An overview of this automatic counting system is provided below.

#### 1.2.2. Outlines for the counting method of the numbers of adult bees and capped brood

This software was developed to count the numbers of adult bees and capped brood accurately. The following explains how to count the number of adult bees present in the image of a comb-frame and how to count the number of capped brood using the image read. These numbers are counted with software using the image from a photo imported into the computer.

The number of adult bees in the colony is obtained by summing the numbers of adult bees on both sides of all the comb-frames with the bees and the number of remaining adult bees in the hive-box without the comb-frame. The number of capped brood in a colony can be obtained by summing the numbers of capped brood on both sides of all the comb-frames without honey bees.

The measurement results obtained by this method, as long as the image remains, can be verified by anyone at any time. The determination procedure of the numbers of adult bees and capped brood is described below.

#### 1.2.3 Determination procedure of the numbers of adult bees and capped brood

STEP-1) The image of comb is binarized, and then the difference in brightness between the adult bee or the capped brood in the comb image and the place where there is no bee, and the features of the adult bee ad the capped brood are used to identify the adult bee and the capped brood, and clarify the counting target. With this identification, you can start the process of automatically counting adult bees or capped brood.
STEP-2) In order to count as accurately as possible, before the binarization process of the image, divide the entire image into several areas that seem to have the same threshold value in each area (up to 4 areas) as below. Then, enter the optimal threshold for each divided area (the maximum number of divided areas 4, the split shape is optional). Examples of division: (1) Distinguish between the capped honey area and the capped brood area. (2) Distinguish between areas where there are few bees and areas where bees are dense. (3) Distinguish between different areas of contrast or lightness (e.g., the part where the light hits and the part of the shadow).
STEP-3) Enter the type of key to mark counted adult bees and capped brood one by one (+, *, numbers, etc.) and the color and size of key ((you can change them after the end of the count).
STEP-4) When the count is performed, while automatically marking those counted adult bee or capped brood in the image, continue counting, the number of counts at that time is displayed at the bottom of the image. After counting the adult bee or capped brood in the image in a short time, the total count in the image is displayed at the bottom of the image. The count status is visually determined, if dissatisfied, by changing the threshold value, recount. In a few tries, even if you change the threshold, if there is no improvement in the count situation, the count of approximate number by automatic measurement in the computer is assumed to have been completed.
STEP-5) After automatic counting by the computer, using the same software, switch to manual count operation, and then correct the count error at the time of automatic counting. Therefore, using a large screen monitor, while further enlarging the image, mark-mistakes of adult bees and capped brood during automatic counting (duplicate count, counting things that are not the object) and forgetting to mark (due to the overlap of bees, the blurred image, the extreme differences in light and dark, etc.) are corrected while visually checking, and all possible efforts are exerted to obtain the number of adult bees or the number of capped brood as accurate as possible (most important and serious work). It should be noted that, even when the software is manual counting, by the removal and grant of the mark, automatically change the number of bees or capped brood. You can resume this fix at any time. If it is determined that the correction is complete, it moves to the measurement of the next new image.
STEP-6) When posting the measured number of adult bees or capped brood in a separate table, call the measured image, check again in the enlarged image, after checking whether there is a count error (correct if you find a mistake), the number is posted to the separate table. The sum total of the numbers of adult bees on all combs and the numbers of adult bees on four walls and bottom in the hive-box is the number of adult bees in a honey bee colony at a certain measurement day, and similarly, the sum total of the numbers of capped brood of all combs is the number of capped brood in a honey bee colony at a certain experiment date.

### 1.3 Supplementary Method S3 (Calculation methods for the intakes of foods and pesticides)

In order to calculate the intakes of food (sugar syrup, pollen paste) and pesticides, let’s explain the calculation procedures using Table S4. Here, it is assumed that the food administration period and the pesticide administration period are the same as the experimental implementation period.

#### 1.3.1 Total number of honey bees during pesticide administration

Here, it is assumed that food and pesticides are continuously administered from the start of the experiment (***ED_0_***) to the end of the experiment (***ED_F_***). We also assume that the larvae are always fed food and pesticides by adult bees, and that the intake of these by drones and queen bee is negligible. In this case, the total number of bees involved in the ingestion of food and pesticides between the start date of the experiment and the end of the experiment can be expressed as the number of capped broods newly fledged and become adult bees plus the number of adult bees that existed on the start date of the experiment.

First, the total number of newly capped brood bees is estimated during the period of administration of food and pesticides (in this example, the experimental period). Suppose that the number of days from the time the pupae in the comb cell is capped to the time they emerge is 12 days. Suppose that the number of days from the time the pupae in the comb cell is capped to the time they emerge is 12 days. Here, consider an experimental date (***ED_i_***) as an example. The number of capped broods on this experimental date (***ED_i_***) is ***CB_i_*** as shown in Table S4. It is assumed that the age-in-days distribution of the number of capped brood on the experimental date ***ED_i_*** is uniform, and that the number of capped broods with each age-in-days of 1 to 12 days is the same for ***CB_i_/***12 (within 12 days until the next experimental date). In other words, in this case, from a certain experiment date (***ED_i_***), capped brood of ***CB_i_***/12 head will continue to fledge every day until the next experiment date (***ED_i+1_***). In addition, even if the number of days until the next experiment date (***ED_i+1_***) exceeds 12 days, it is assumed that ***CB_i_***/12 adult bee will continue to newly emerge. The number (***NB_i_***) in which capped brood will fledge and new adult bees will emerge from a certain experimental date (***ED_i_***) to the next date (***ED_i+1_***) is obtained by the following equation.

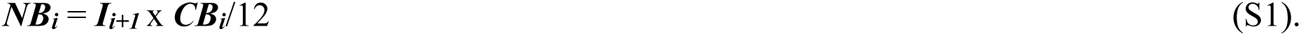

Here, ***I_i_*** is up to ***I_F_***.

Since food (sugar syrup, pollen paste) and pesticides are administered from the start of the experiment date (***D_0_***) to the date of the end of the experiment (***D_F_***), the total number (***C_T_***) of capped broods that fledge and new adult bees emerge in the experimental period (***I_T_***) is obtained by the following equation.

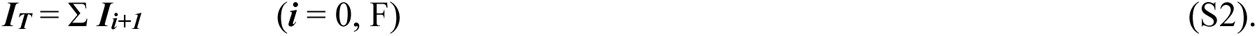

Therefore, from the start of the experiment (***ED_0_***) to the date of the end of the experiment (***ED_F_***), the total number of adult bees (***AB_T_***) involved in the ingestion of food and pesticides is given by adding the number of adult bees (***AB_0_***) present at the start of the experiment and the number of capped brood at the end of the experiment to the total number of newly adult bees (***NB_T_***) that have emerged from the pupa. Because capped brood at the end of experiment would have already ingested the pesticide-containing food by the end of experiment.

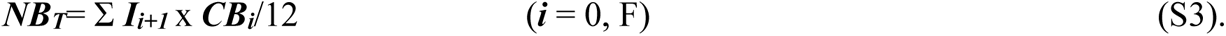

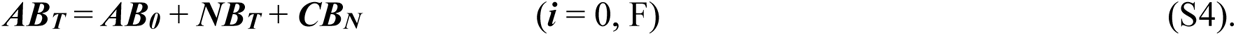

Here, ***t_i_*** is up to ***t_F_***.

#### 1.3.2 Total intake of pesticide or food per colony during pesticide administration

Consider that the intakes of food (sugar syrup, pollen paste) and pesticides taken by one bee colony are equal to their consumptions measured on each experimental date, respectively. In addition, the intake of pesticides is calculated from the consumption of food containing a pesticide. That is, the consumptions (***SS_T_***) of sugar syrup, pollen paste (***PP_T_***) and pesticide (***P_T_***) by the bee colony are each obtained by the following equations.

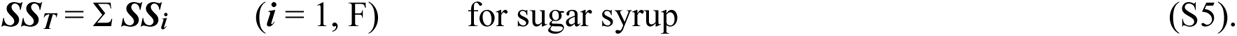

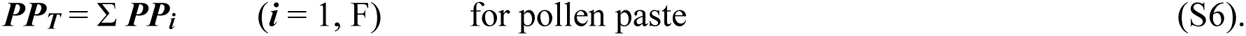

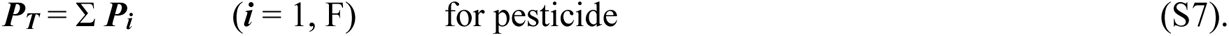

#### 1.3.3 Total intake of pesticide or food per bee during pesticide administration

During the pesticide administration period (here, the same as the experimental period), the total intakes of sugar syrup (***SS_B_***), pollen paste (***PP_B_***), and pesticides (***P_B_***) ingested by one bee are determined by the following equations, respectively.

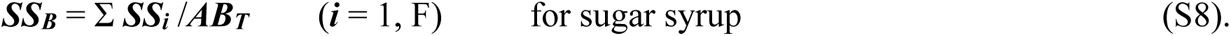

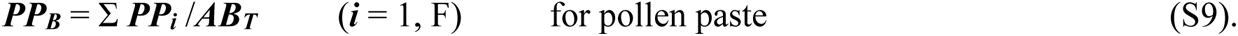

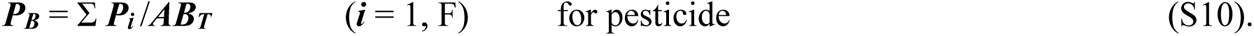

#### 1.3.4 Average daily intake of pesticide or food per colony during pesticide administration

The average intake of sugar syrup (***SS_D_***), pollen paste (***PP_D_***), and pesticide (***P_D_***) ingested by bee colonies in one day during the pesticide administration period is calculated by the following equations, respectively.

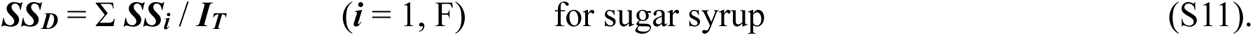

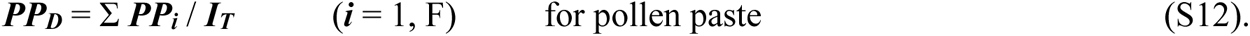

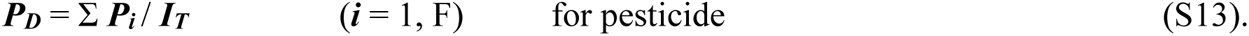

Where t_T_ denotes the total number of days from the start of experiment to the end of experiment, which is equal to the administration period of pesticide.

#### 1.3.5 Average daily intake of pesticide or food per bee during pesticide administration

The average intake of sugar syrup (***SS_BD_***), pollen paste (***PP_BD_***), and pesticide (***P_BD_***) ingested by one bee in one day during the pesticide administration period is determined by the following equations.

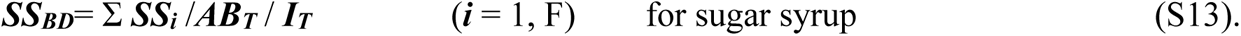

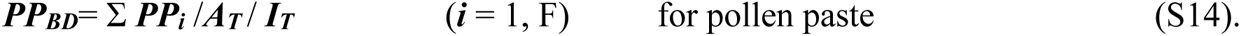

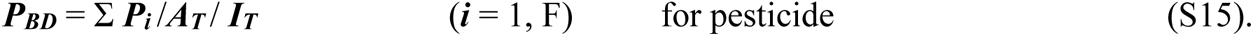

## 2 Supplementary Figures (Figures. S1 to S7)

**Supplementary Figure S1.**
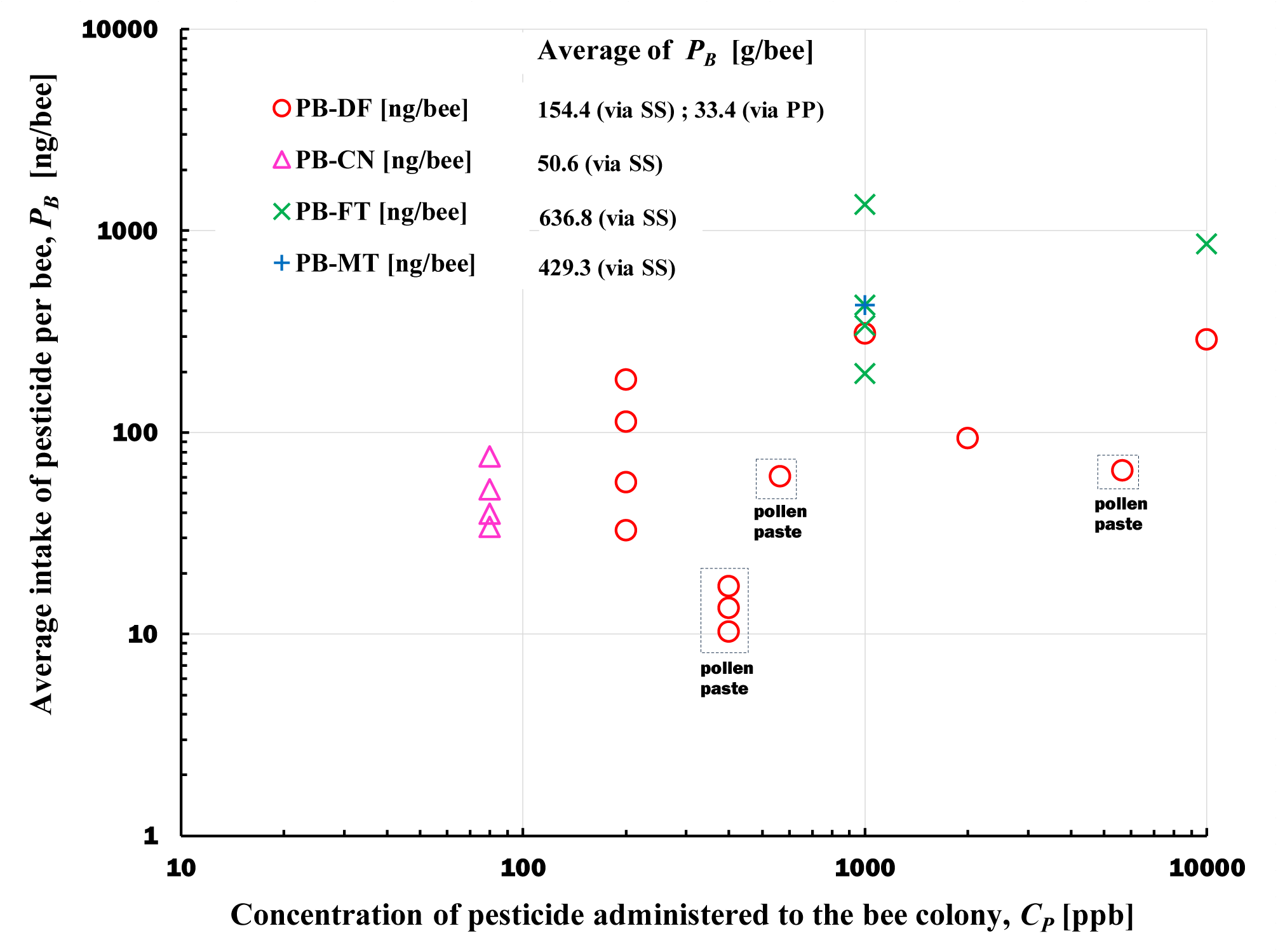
Relationship between the concentration of pesticide administered to the bee colony (*C_P_*) and the total intake of pesticide per bee (***P_B_***) during pesticide administration. **P_B_:** Total intake of a pesticide ingested by one bee during the administration period. **P_B-DF_:** Total intake of dinotefuran ingested by one bee during the administration period. **P_B-CN_:** Total intake of clothianidin ingested by one bee during the administration period. **P_B-FT_:** Total intake of fenitrothion ingested by one bee during the administration period. **P_B-MT_:** Total intake of malathion ingested by one bee during the administration period. The data surrounded by a dotted square is the ***P_B_*** value where the pesticide was administered to the bee colony via pollen paste, and the rest is the ***P_B_*** value where the pesticide was administered via sugar syrup.

**Supplementary Figure S2.**
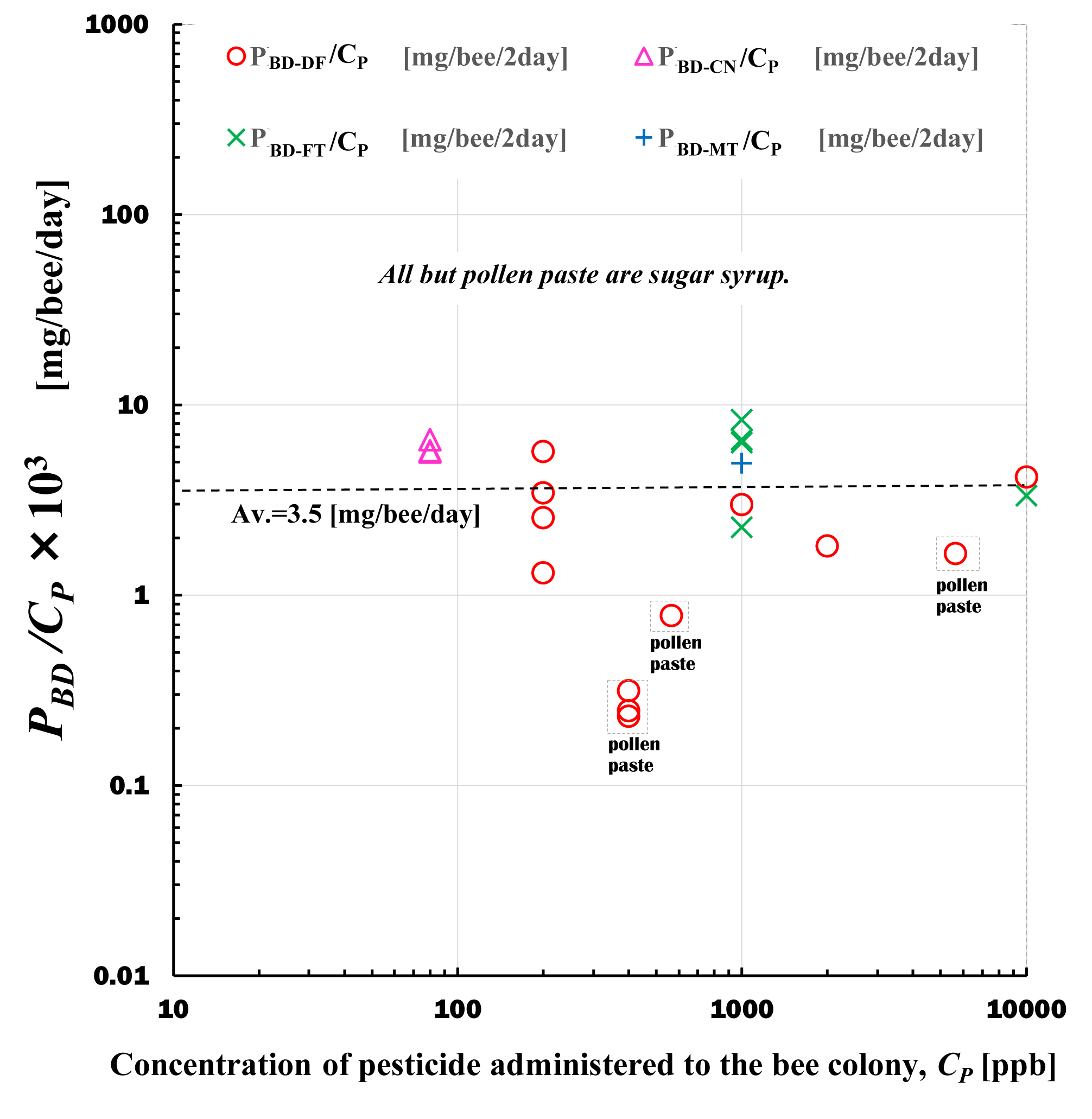
Ratio of the average daily intake of pesticide per bee (***P_BD_***) during the pesticide administration period to the concentration of pesticide administered to the bee colony (***C_P_***). In the figure, the data surrounded by a dotted square is the ***P_BD_***/***C_P_*** value where the pesticide was administered to the bee colony via pollen paste, and the rest is the ***P_BD_***/***C_P_*** value where the pesticide was administered via sugar syrup. The data surrounded by a dotted line in the upper rectangle labeled pollen paste in the figure shows the daily intake of pesticides by one bee when “pollen paste” containing pesticides is administered to a bee colony as a vehicle. All other data show the daily intake of pesticides per bee when pesticides are administered using sugar syrup as the vehicle.

**Supplementary Figure S3.**
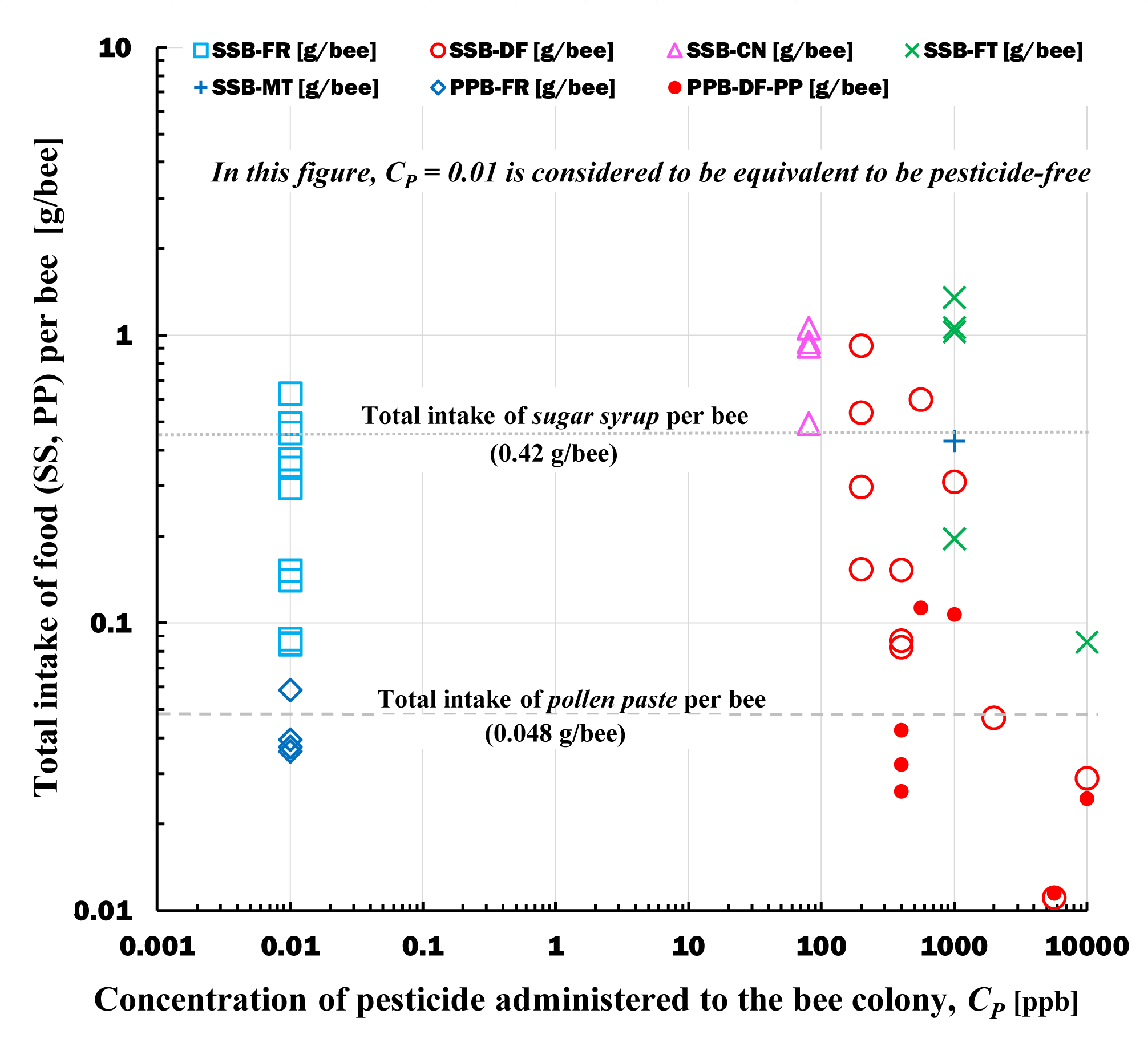
Relationship between the administration-concentration of pesticide to the bee colony and the total intake of food (sugar syrup, pollen paste) per bee during the administration period. SS: sugar syrup, PP: pollen paste, FD: food (SS, PP). **SS_B-FR_:** Total intake of pesticide-free SS per bee during the administration period. **SS_B-DF_:** Total intake of SS containing dinotefuran per bee during the administration period. **SS_B-CN_:** Total intake of SS containing clothianidin per bee during the administration period. **SS_B-FT_:** Total intake of SS containing fenitrothion per bee during the administration period. **SS_B-FT_:** Total intake of SS containing malathion per bee during the administration period. **PP_B-FR_:** Total intake of pesticide-free PP per bee during the administration period. **PP_B-DF_:** Total intake of PP containing dinotefuran per bee during the administration period.

**Supplementary Figure S4.**
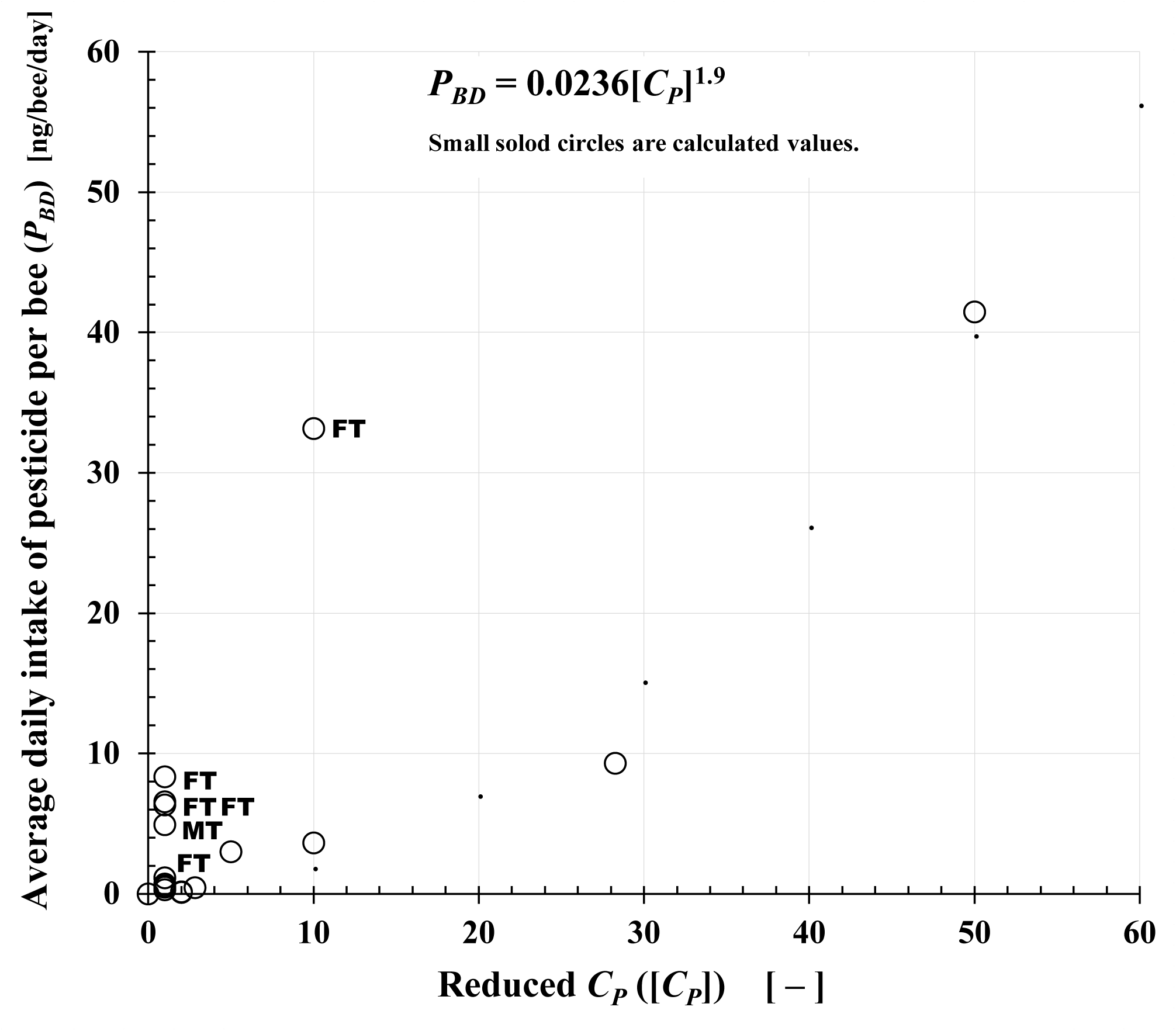
Relationship between the reduced concentration of pesticide administered to the bee colony, [***C_P_***], and the average daily intake of pesticide per bee per day (***P_BD_***) during pesticide administration. The reduced administration-concentration [***C_P_***] is a dimensionless concentration expressed by dividing the insecticide administration-concentration to a bee colony by each (as a reference concentration) of dinotefuran (DF) 0.2 ppm, clothianidin (CN) 0.08 ppm, fenitrothion (FT) 1 ppm, and malathion (MT) 1 ppm, which have almost the same insecticidal activity (acute toxicity) against stink bugs.

**Supplementary Figure S5.**
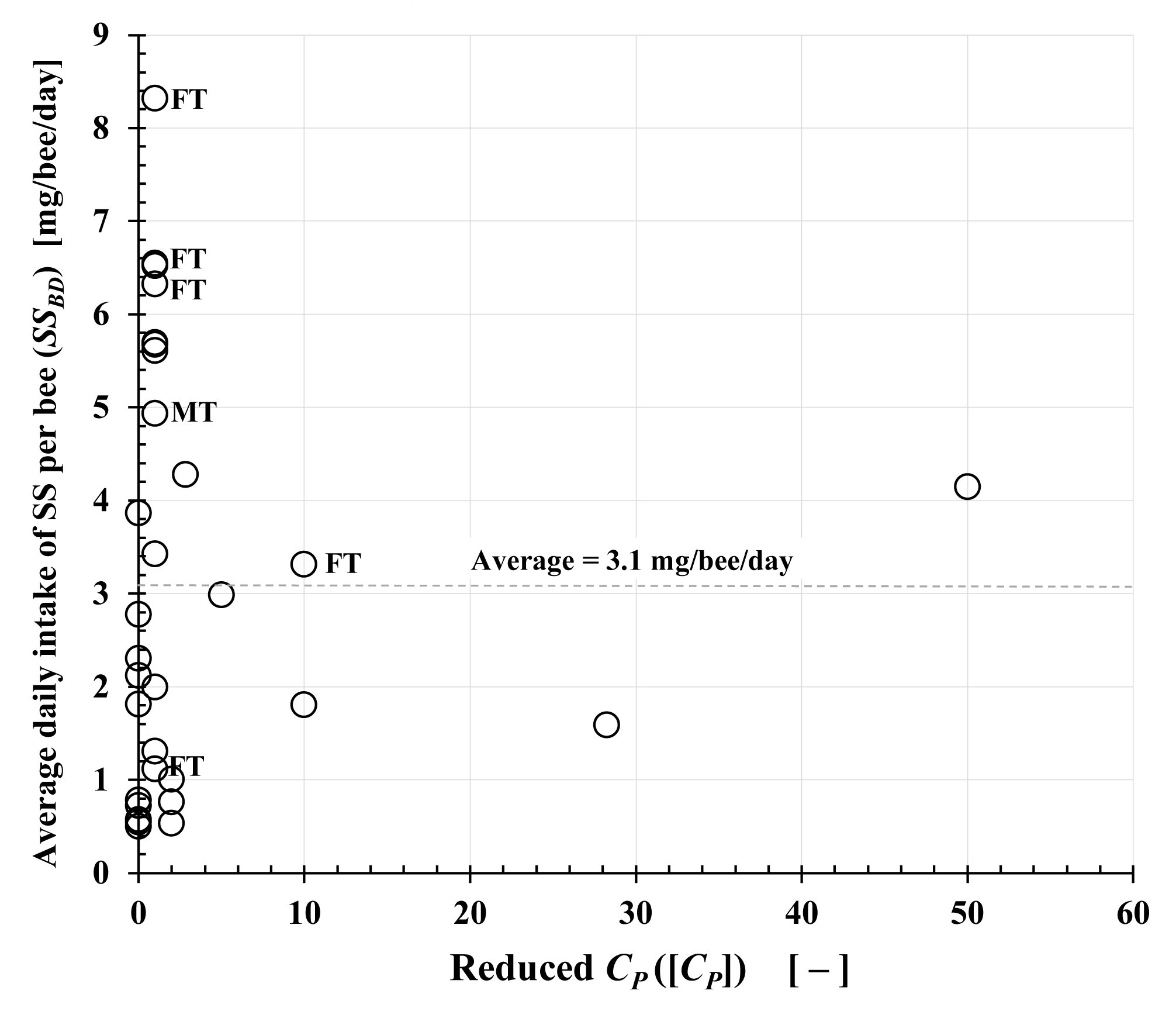
Relationship between the reduced concentration of pesticide administered to the bee colony, [***C_P_***], and the average intake of sugar syrup per bee per day (***SS_BD_***) during pesticide administration. The reduced administration-concentration [***C_P_***] is a dimensionless concentration expressed by dividing the insecticide administration-concentration to a bee colony by each (as a reference concentration) of dinotefuran (DF) 0.2 ppm, clothianidin (CN) 0.08 ppm, fenitrothion (FT) 1 ppm, and malathion (MT) 1 ppm, which have almost the same insecticidal activity (acute toxicity) against stink bugs.

**Supplementary Figure S6.**
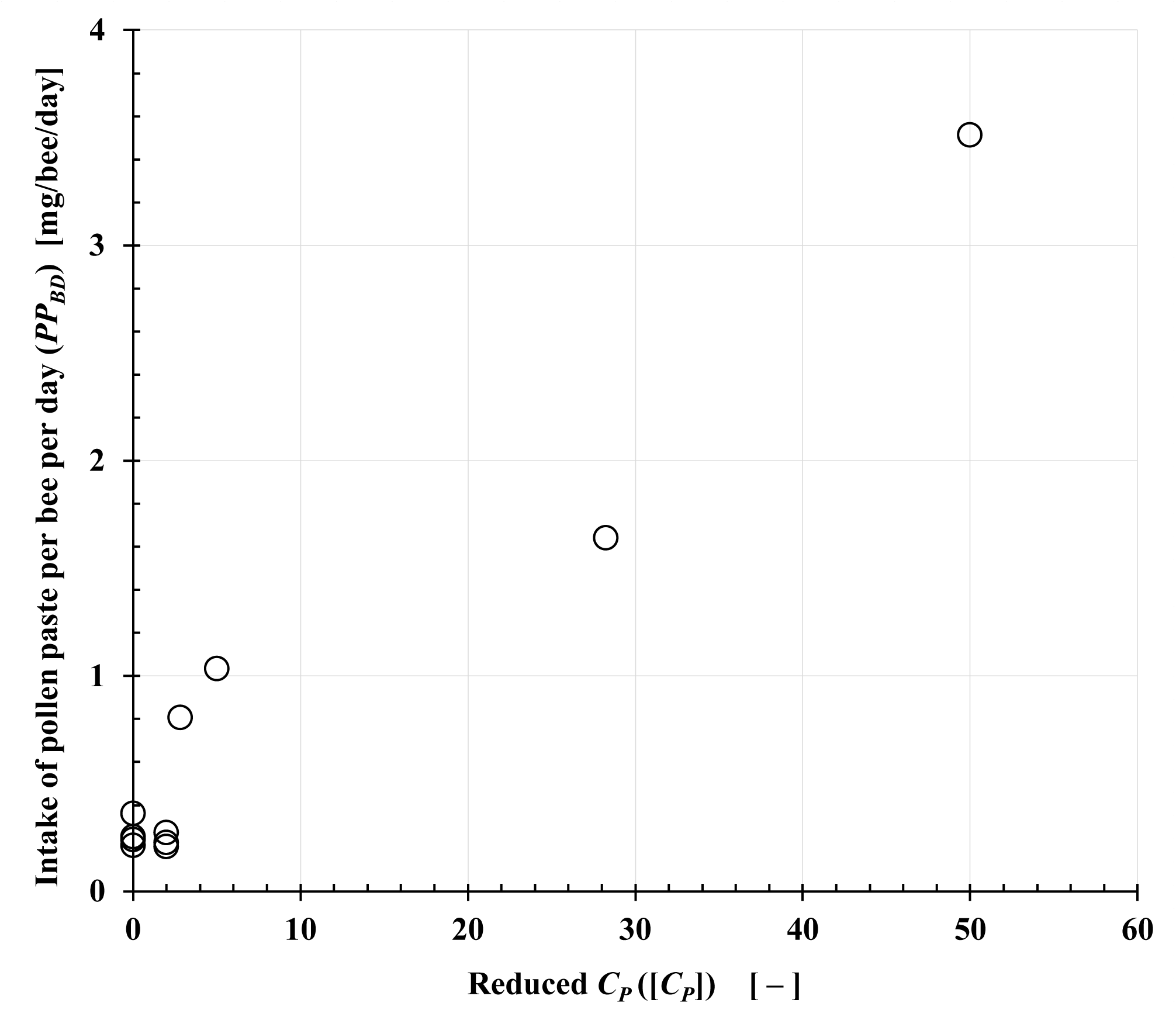
Relationship between the reduced concentration of pesticide administered to the bee colony, [***C_P_***], and the average intake of pollen paste per bee per day (***PP_BD_***) during pesticide administration. The reduced administration-concentration [*C_P_*] is a dimensionless concentration expressed by dividing the insecticide administration-concentration to a bee colony by each (as a reference concentration) of dinotefuran (DF) 0.2 ppm, clothianidin (CN) 0.08 ppm, fenitrothion (FT) 1 ppm, and malathion (MT) 1 ppm, which have almost the same insecticidal activity (acute toxicity) against stink bugs.

**Supplementary Figure S7.**
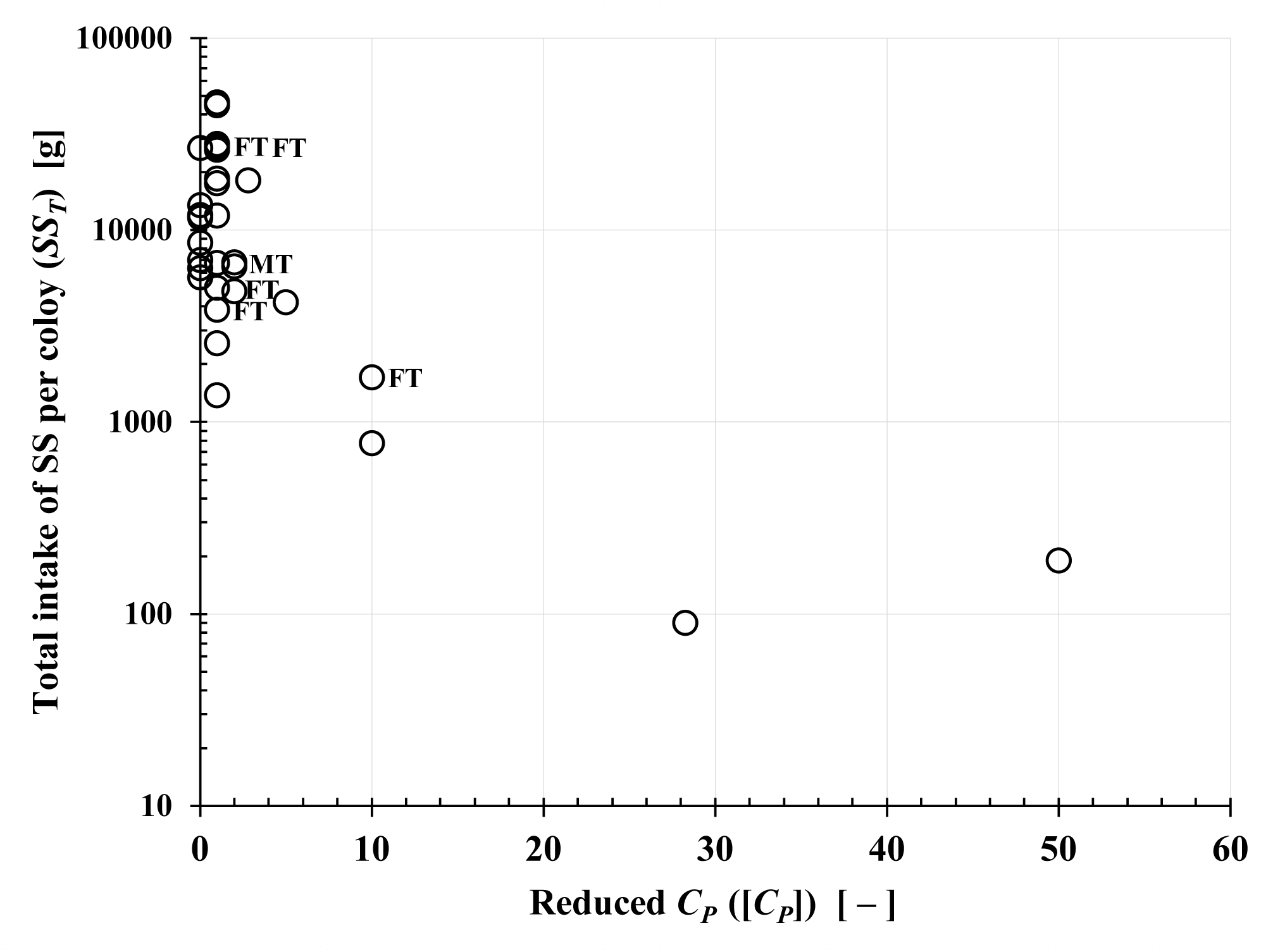
Relationship between the reduced concentration of pesticide administered to the bee colony, [***C_P_***], and the total intake of sugar syrup per colony (***SS_T_***) during pesticide administration. The reduced administration-concentration [***C_P_***] is a dimensionless concentration expressed by dividing the insecticide administration-concentration to a bee colony by each (as a reference concentration) of dinotefuran (DF) 0.2 ppm, clothianidin (CN) 0.08 ppm, fenitrothion (FT) 1 ppm, and malathion (MT) 1 ppm, which have almost the same insecticidal activity (acute toxicity) against stink bugs.

## 3 Supplementary Tables (Tables. S1 to S5)

**Supplementary Table S1.**
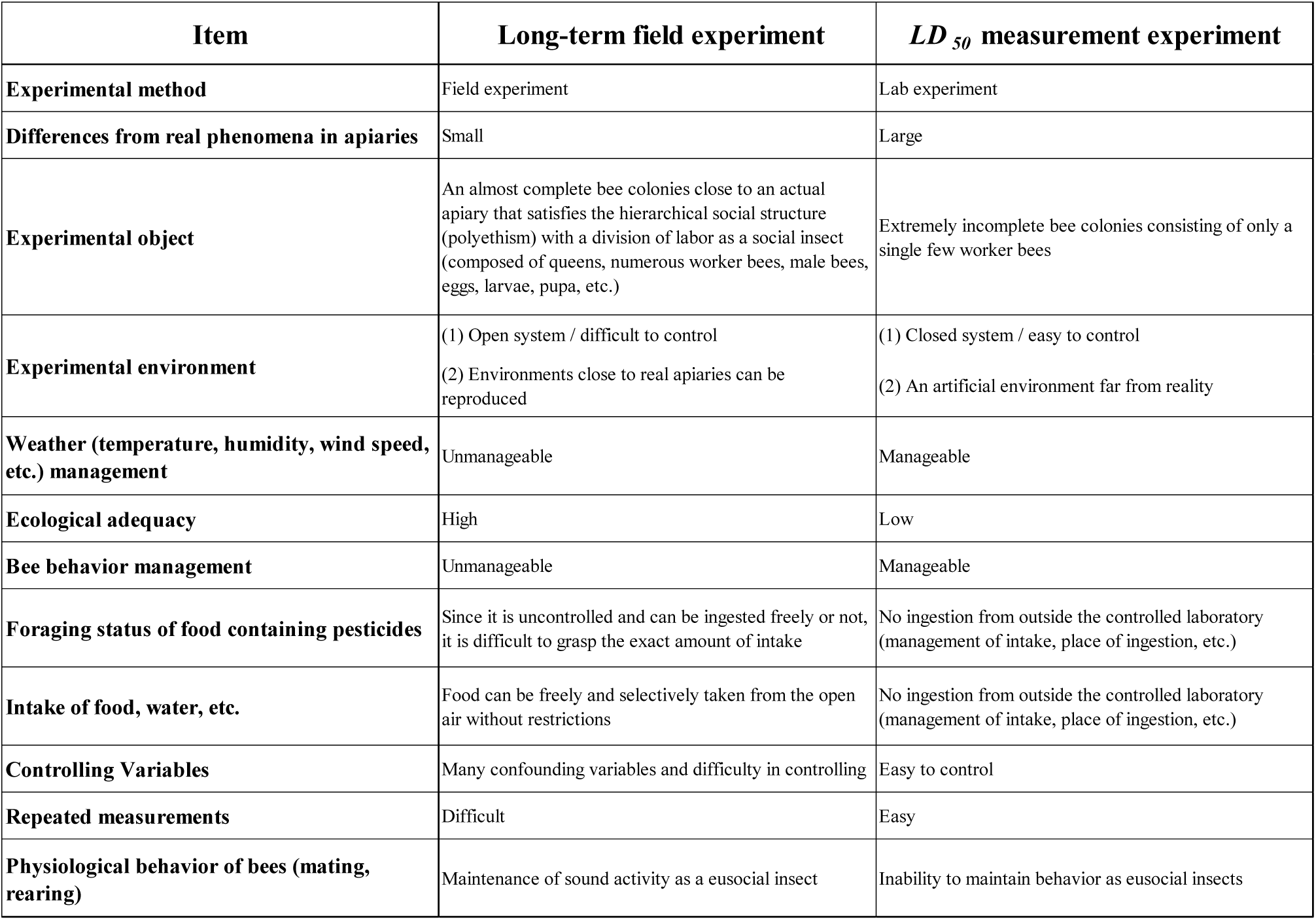
Comparison between long-term field experiments and *LD_50_* measurement experiments in the investigation of the effects of pesticides on bees.

**Supplementary Table S2.**
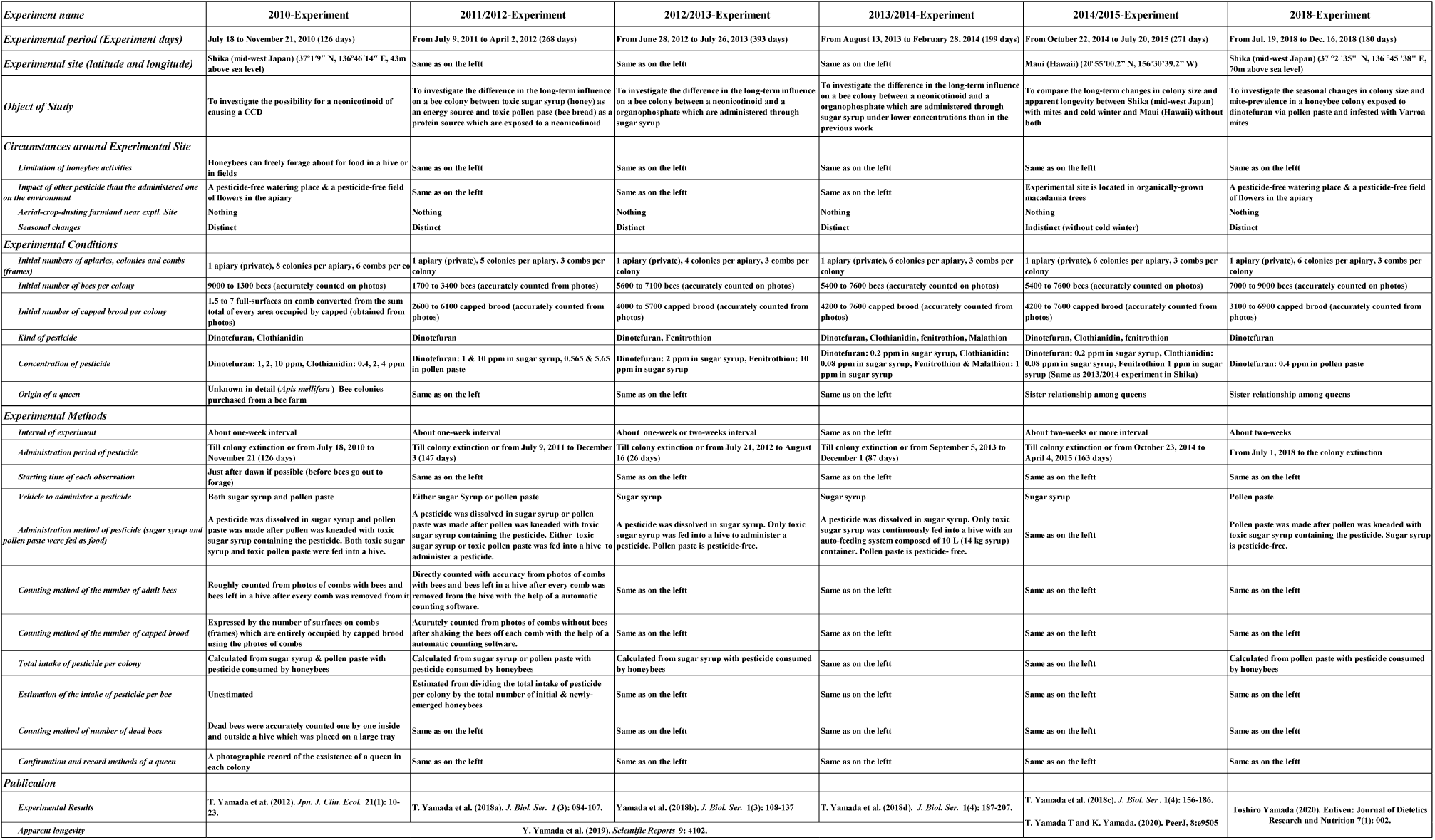
Experimental conditions of six long-term field experiments for honeybee colonies.

**Supplementary Table S3.**
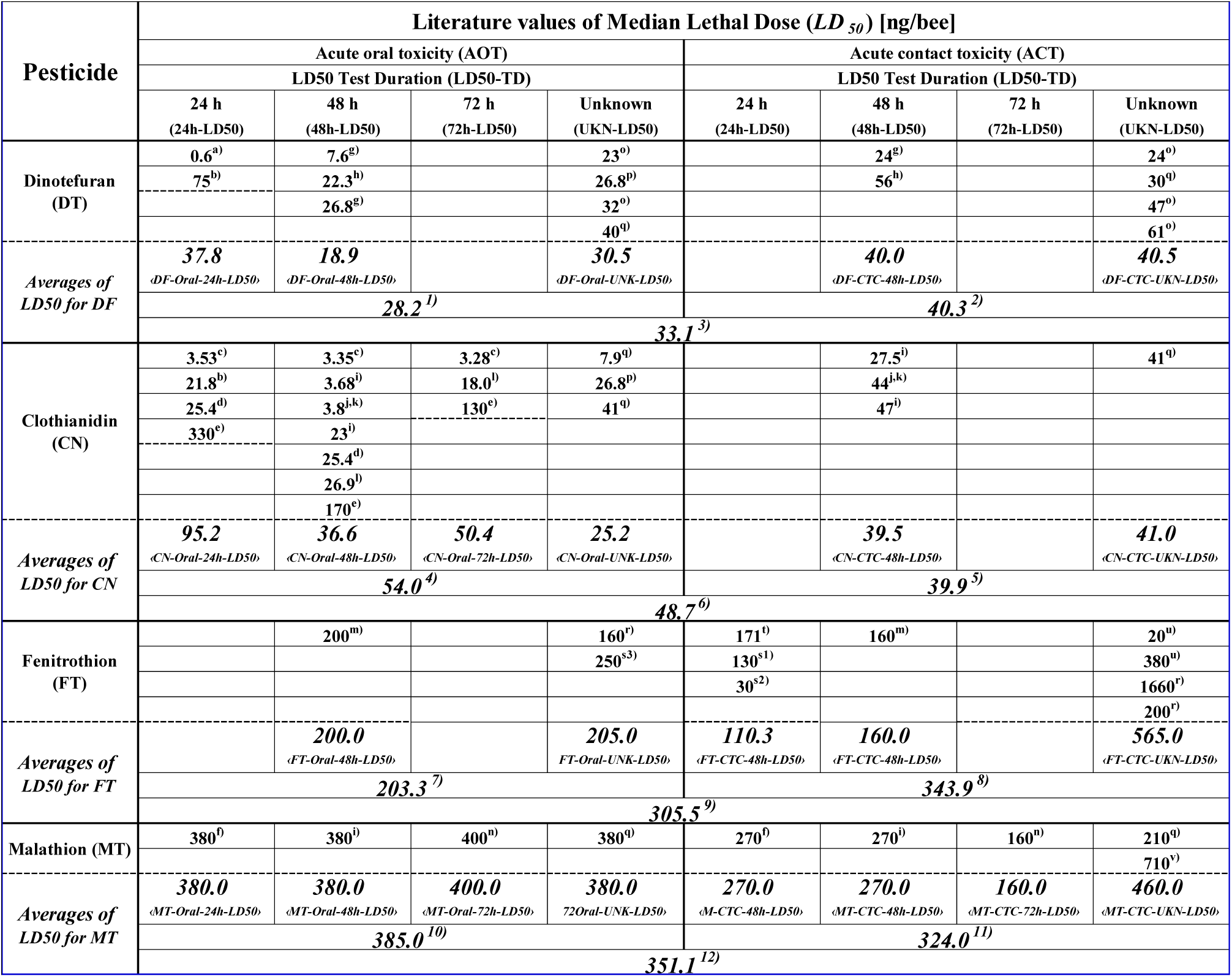
Literature values of median lethal doses (***LD_50_***) for the honeybee and various average values of the ***LD_50_*** data for dinotefuran, clothianidin, fenitrothion and malathion. **^1)^** Average of all ***LD_50_*** data on acute oral toxicity of dinotefuran (〈D F-Oral-LD50〉). **^2)^** Average of all ***LD_50_*** data on acute contact toxicity of dinotefuran (〈D F-CTC-LD50〉). **^3)^** Average of all ***LD_50_*** data on acute oral and contact toxicities of dinotefuran (〈DF-LD50〉) **^4)^** Average of all ***LD_50_*** data on acute oral toxicity of clothianidin (〈CN-Oral-LD50〉). **^5)^** Average of all ***LD_50_*** data on acute contact toxicity of clothianidin (〈CN – CTC –LD50〉). **^6)^** Average of all ***LD_50_*** data on acute oral and contact toxicities of clothianidin (〈CN – LD50〉) **^7)^** Average of all ***LD_50_*** data on acute oral toxicity of fenitrothion (〈FT-Oral-LD50〉). **^8)^** Average of all ***LD_50_*** data on acute contact toxicity of fenitrothion (FT – CTC –LD50). **^9)^** Average of all ***LD_50_*** data on acute oral and contact toxicities of fenitrothion (〈FT – LD50〉) **^10)^** Average of all ***LD_50_*** data on acute oral toxicity of malathion (〈MT-Oral-LD50〉). **^11)^** Average of all ***LD_50_*** data on acute contact toxicity of malathion (〈MT – CTC –LD50〉). **^12)^** Average of all ***LD_50_*** data on acute oral and contact toxicities of malathion (〈MT – LD50〉) The list of literatures cited is shown in Supplementary Table S3a.

**Supplementary Table S3a.**
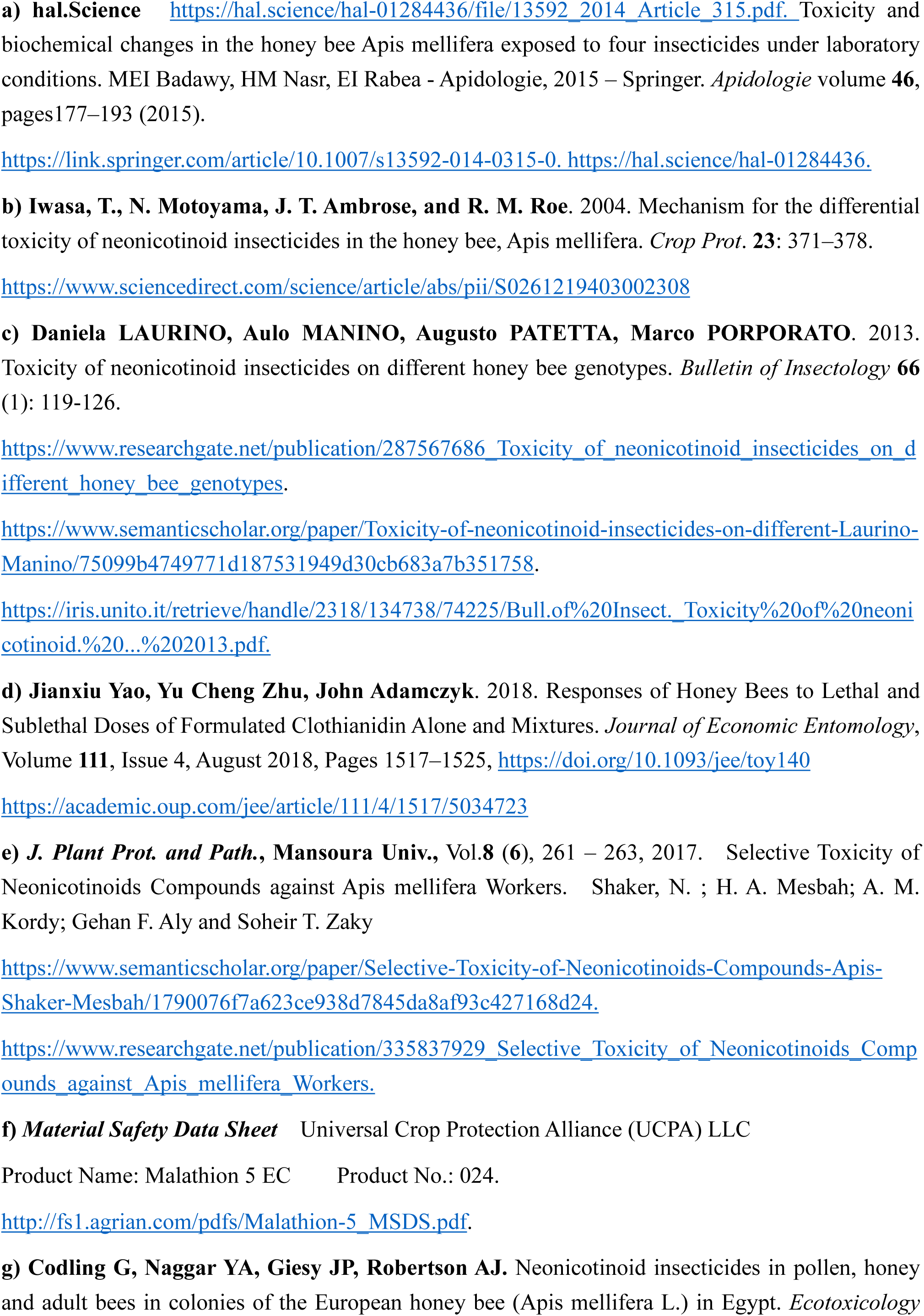
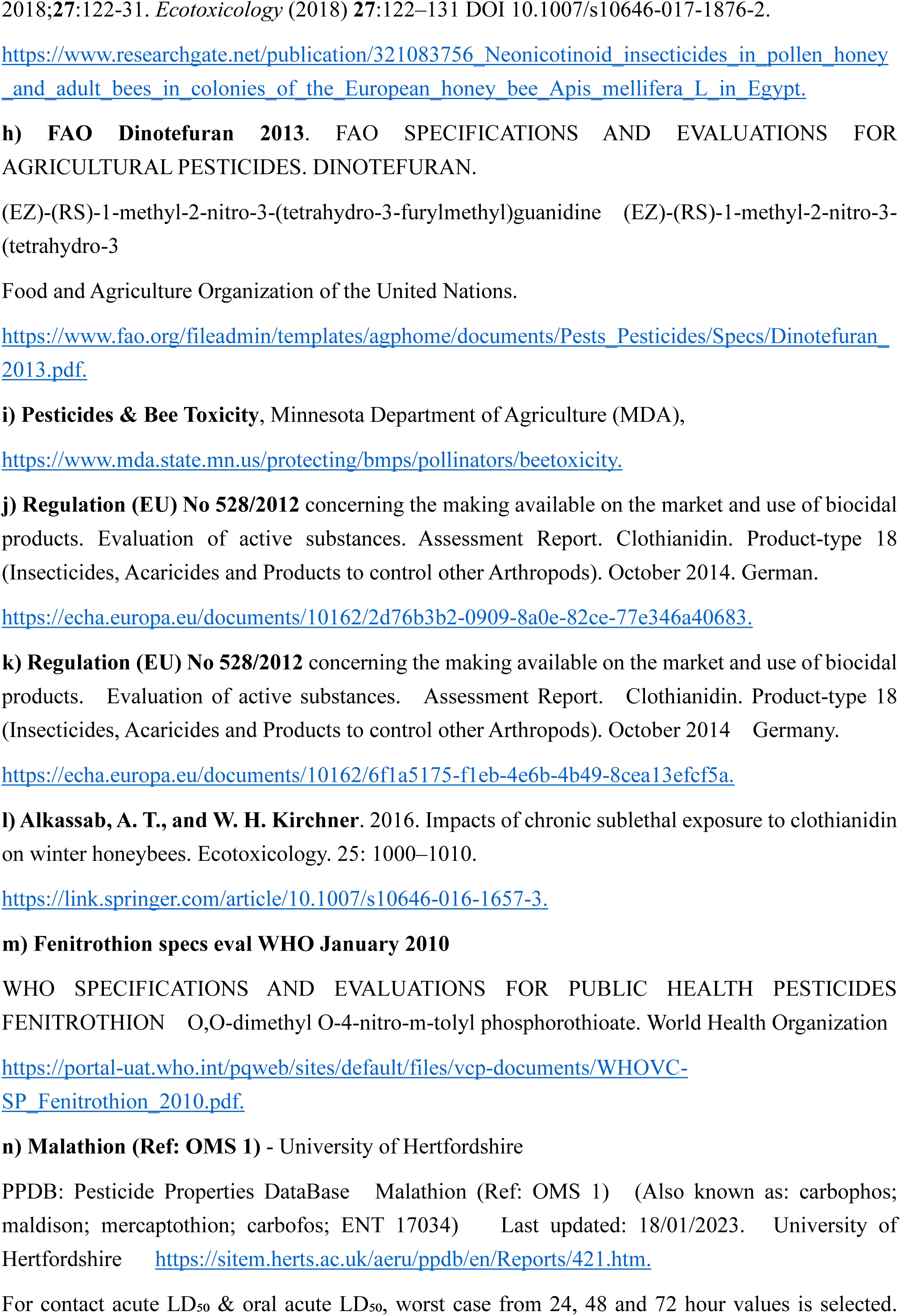
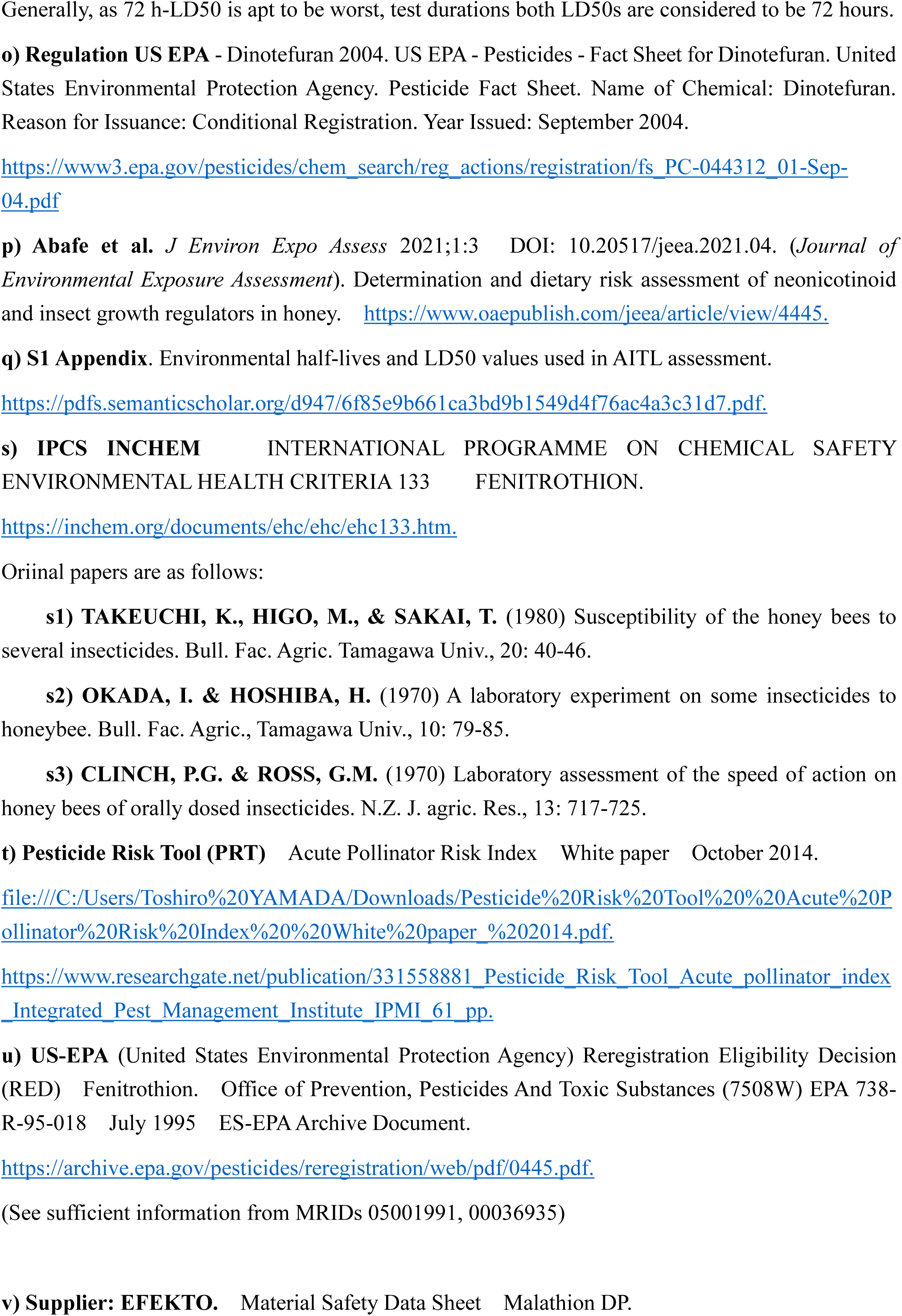
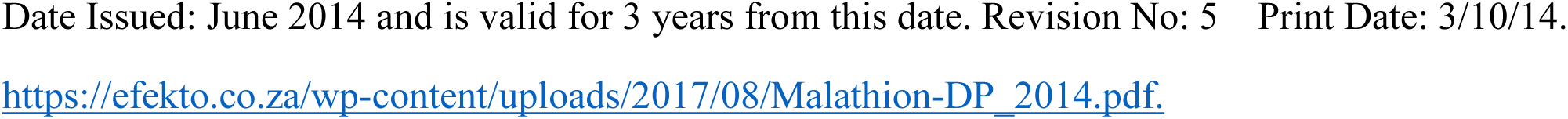
List of literatures cited in Supplementary Table S3.

**Supplementary Table S4.** Example data for intakes of food and pesticide. Interval =The interval between two adjacent experimental dates (***I_i_***); ***I_i_*** = ***ED_i_*** – ***ED_i-1_*** Adult: The number of adult bees; Brood: The number of capped brood. Total number of experiment days from ***ED_0_*** till ***ED_F_*** (***t_T_***): ***I_T_* = Σ*I_i_*** (***i*** =1, F) Total consumption of sugar syrup from ***ED_0_*** till ***ED_F_*** (***SS_T_***): ***SS_T_* = Σ*SS_i_*** (***i*** =1, F) Total consumption of pollen paste from ***ED_0_*** till ***ED_F_*** (***PP_T_***): ***PP_T_* = Σ*PP_i_*** (***i*** =1, F) Total consumption of pesticide from ***ED_0_*** till ***ED_F_*** (***P_T_***): ***P_T_* = Σ*PP_i_*** (***i*** =1, F)

| Exptl. Date | Interval | Adult Bee | Capped Brood | Sugar Syrup | Pollen Paste | Pesticide |
| --- | --- | --- | --- | --- | --- | --- |
|  | [day] | [head] | [head] | [g] | [g] | [μg] |
| $ED_0$ | 0 | $AB_0$ | $CB_0$ | 0 | 0 | 0 |
| $ED_1$ | $I_1$ | $AB_1$ | $CB_1$ | $SS_1$ | $PP_1$ | $P_1$ |
| $ED_2$ | $I_2$ | $AB_2$ | $CB_2$ | $SS_2$ | $PP_2$ | $P_2$ |
| $ED_3$ | $I_3$ | $AB_3$ | $CB_3$ | $SS_3$ | $PP_3$ | $P_3$ |
| $ED_4$ | $I_4$ | $AB_4$ | $CB_4$ | $SS_4$ | $PP_4$ | $P_4$ |
| $ED_5$ | $I_5$ | $AB_5$ | $CB_5$ | $SS_5$ | $PP_5$ | $P_5$ |
| $ED_6$ | $I_6$ | $AB_6$ | $CB_6$ | $SS_6$ | $PP_6$ | $P_6$ |
| $ED_F$ | $I_F$ | $AB_F$ | $CB_F$ | $SS_F$ | $PP_F$ | $P_F$ |

**Table S5.** Each average value of ***SS_BD_***, ***PP_BD_*** and ***P_BD_*** and that of ***SS_B_***, ***PP_B_*** and ***P_B_*** during the pesticide administration period. DF denotes dinotefuran, CN denotes clothianidin, FT denotes fenitrothion, and MT denotes Malathion. SS denotes sugar syrup and PP denotes Pollen Paste. For ***SS_BD_***, ***PP_BD_***, ***P_BD_***, ***SS_B_***, ***PP_B_*** and ***P_B_*** see Supplementary Method S3.

| Target Data for Average Value | Vehicle to administer a pesticide | Total intake per bee |  |  | Average daily intake per bee |  |  |
| --- | --- | --- | --- | --- | --- | --- | --- |
| | | Sugar Syrup ($SS_B$) | Pollen Paste ($PP_B$) | Pesticide ($P_B$) | Sugar Syrup ($SS_{BD}$) | Pollen Paste ($PP_{BD}$) | Pesticide ($P_{BD}$) |
|  |  | [g/bee] | [g/bee] | [ng/bee] | [mg/bee/day] | [mg/bee/day] | [ng/bee/day] |
| All data | — | 0.4236 | 0.0481 | — | 3.1160 | 0.7997 | — |
| Pesticide-free data | — | 0.2863 | 0.0429 | — | 1.6666 | 0.2694 | — |
| All pesticide data | — | 0.4922 | 0.0511 | 295.3 | 3.8407 | 1.1027 | 5.5956 |
| DF dosage data | SS+PP | 0.2699 | 0.0511 | 106.6 | 2.4620 | 1.1027 | 5.0608 |
|  | SS | 0.3293 | 0.0662 | 158.3 | 3.0504 | 2.2752 | 7.2395 |
|  | PP | 0.1867 | 0.0451 | 34.3 | 1.6364 | 0.6337 | 2.0106 |
| CN dosage data | SS | 0.8559 | — | 68.5 | 6.0954 | — | 0.4876 |
| FT dosage data | SS | 0.7477 | — | 902.9 | 5.3559 | — | 11.3267 |
| MT dosage data | SS | 0.4293 | — | 429.3 | 4.9344 | — | 4.9344 |

